# Unraveling the potential of short and long read sequencing for human genome profiling

**DOI:** 10.64898/2026.04.20.719568

**Authors:** Aurélie Leduc, Asmae Bachr, Florian Sandron, Marc Delépine, Damien Delafoy, Cédric Fund, Christian Daviaud, Stéphane Meslage, Violette Turon, Delphine Bacq-Daian, Francis Rousseau, Robert Olaso, Jean-François Deleuze, Zuzana Gerber, Vincent Meyer

**Author notes:** These authors contributed equally and share first authorship (Aurélie Leduc, Asmae Bachr). These authors co-supervised the work (Zuzana Gerber, Vincent Meyer).

## Abstract

**Background:** Short read sequencing technologies have dominated the field of human whole genome sequencing in the past years in terms of cost, throughput, and accuracy. However, thanks to recent technological evolution, long read approaches have become increasingly competitive and complementary to short reads. With the gap in the cost per genome closing slowly between both approaches, long reads might replace short read sequencing in future research and clinical applications. Still, comprehensive evaluation is necessary to conclude on the performance and general advantages of each technology.

**Results:** In this study, we compared the latest chemistries of major suppliers of short and long read technologies: Illumina short reads, Illumina Complete Long Reads (ICLR), Pacific Biosciences HiFi reads (PacBio), and Oxford Nanopore Technologies long reads (ONT). Using the HG002 human reference sample and established bioinformatics guidelines, we assessed their variant calling performance against the latest available truth sets at different levels of coverage. For single nucleotide variant detection, all technologies were equivalent. Despite the latest improvements in chemistry, indel calling with ONT continues to lag in accuracy behind other technologies. In contrast, long reads delivered a clear advantage in structural variant detection, surpassing short reads in both accuracy and sensitivity. The hybrid ICLR approach achieved intermediate performance, narrowing the gap between short and long read sequencing. Furthermore, long reads enhanced haplotype-phasing resolution, enabling the phasing of over 80% of the genome.

**Conclusions:** These findings highlight the specific strengths and limitations of recent sequencing technologies, aiding the decision-making in future research projects, technological platforms development, and clinical applications.

## 1 Background

Supported by the fast evolution of sequencing technologies nowadays, human genomics progressively integrates research and clinical applications [1, 2]. A range of solutions is now available to fit the requirements in diverse fields of research. Sequencing enables genetic profiling, expression, methylation [3, 4], DNA conformation and metagenomics analyses [5] in diverse experimental contexts.

Short read technologies offer robust performance, the highest read quality, and the lowest cost-per-genome at present, but with a maximum read length of only 600 base pairs (bp). While Illumina remains the most established sequencing technology on the market today, other providers, such as MGI [6] and Ultima Genomics [7] propose competitive offers with high-throughput, cost-efficient strategies and data quality similar to Illumina. Thanks to recent improvements in long read technologies, represented by Pacific Biosciences (PacBio) and Oxford Nanopore Technologies (ONT), long read approach becomes increasingly competitive on the sequencing market and compatible with human genomics research and clinical applications [8, 9]. This approach enables access to complex regions of the genome, so far unexplored [10], and brings new insights into the structural organization of the genome at individual, population [11, 12], or pangenomic level [13]. Recent technological innovations include improved read quality, increased throughput capacity, more consistent results, and decreased cost-per-genome, all while providing read lengths far above 10 kilobases (kb) long.

To address the increasing demand for long read data, Illumina developed a hybrid strategy called Illumina Complete Long Reads (ICLR) based on bioinformatics reconstruction of long fragments from short read data. This approach allows Illumina to provide haplotype information for improved single nucleotide variant (SNV), small insertion-deletions (indel), and structural variant (SV) calling, as well as haplotype phasing with no dedicated long read sequencer.

ONT technology relies on nanopores: DNA molecule passing through a nanopore causes a sequence-specific disturbance in the electric current, which is detected and converted into nucleotide sequence using a machine learning algorithm. ONT improves continually the sequencing chemistry, nanopore architecture, as well as the modelisation of the signal properties and correction. This technology can provide the longest reads with Q20+ quality. With the current V14 chemistry, the recommended read length with ONT standard approach is 10 – 30 kb, which provides the highest sequencing yield. ONT PromethION sequencer can provide 90 – 120 Gb yield per flow cell, equivalent to a 30 – 40× depth of coverage of the human genome according to ONT specifications. Alternatively, the read length can reach several megabases with ONT ultra-long approach [14], though the sequencing yield is lower.

Single Molecule Real-Time sequencing technology of PacBio relies on a modified polymerase to sequence DNA fragments using tagged nucleotides and fluorescence detection. The PacBio technology is based on circular consensus sequencing [15]. It enables efficient error correction using machine learning algorithm, providing high sequencing quality (Q30+), which is however limited to median read length not exceeding 25 kb. PacBio Revio sequencer generates on average 30× – 40× depth of coverage of the human genome per SMRT Cell according to chemistry version.

Currently, each of the technologies has specific strengths and weaknesses, impacting the quality and precision of the information produced. The most common application in human genomics is the profiling of genetic variants including SNVs, indels, and SVs. The field of genomics is very dynamic and while variant calling relies on well established general principles, new algorithms and methods are under continuous development. New or upgraded tools appear frequently, which require new benchmarks for further integration into data processing workflows [16–18].

In the context of human genome studies, mapping-based approaches that align reads to the human reference genome [19, 20] are widely employed for both short and long read sequencing technologies. The most common tools to process large amount of sequences are based on k-mer indexing combined to Burrows-Wheeler transformation incorporating various adaptations to manage mapping accuracy [21, 22]. The alignments are then processed through variant caller programs to identify differences between the piled-up reads and the selected reference [23]. For short reads, the GATK toolkit [24] has been established as a widely used solution for SNV and indel calling, accompanied by additional solutions sharing similar principles [25]. More recently, emerging approaches that exploit basecalling and mapping information within deeplearning models, notably represented by DeepVariant and Clair3 programs, improved the accuracy and precision [26, 27]. This generation of callers is also available for long reads, with dedicated models addressing ONT- and PacBio-specific data properties. Besides that, the best practices have been evolving recently towards the use of pangenome references and graph-genome mapping algorithms. Specific tools for short reads pangenome analysis are available and evolving, such as vg and vg giraffe [28, 29]. Furthermore, Illumina provides a dedicated process within their commercial solution DRAGEN [30]. This new paradigm is also adapted and progressively migrating to long reads sequencing studies [31, 32].

SV calling from short reads is possible by leveraging paired-end alignment information and depth of coverage, which broadly covers the different SV types using Delly [33], Lumpy [34], GRIDSS [35], Manta [36] or GATK [37]. Tools exist for specific SV types such as mobile elements, local pattern expansions or large Copy Number Variations [38]. In spite of a multitude of tools, SV calling from short read data is challenging, even with a number of complementary technologies such as CGH-array [39], barcoding-DNA or linked-reads [40, 41]. Owing to the capacity of long reads to provide direct and comprehensive reads, sometimes spanning the whole length of complex and repetitive regions that are inaccessible with short reads, long read sequencing represents a breakthrough in SV profiling [42–44] and it is consequently considered the Method of the Year 2022 [45]. Both PacBio and ONT provide a dedicated bioinformatics workflow for data processing. These workflows leverage the strengths and mitigate the weaknesses specific to each technology. In the absence of paired-end data, SV calling from long reads relies on local alignment optimization and identification of specific features of different SV types. The performance of SV calling improves continually with new software updates and new tools [10, 46–48].

In this study, in a context where the technological and experimental options for clinical research are continually evolving, we present an evaluation of state-of-theart sequencing technologies for human genome profiling, analyzed with gold standard bioinformatics tools, against high-confidence truth sets for the reference sample HG002. Compared to previous benchmarks [49–51], we present a comprehensive comparison among four cutting-edge sequencing technologies. Using the latest chemistry and the latest sequencer generations for each technology, we provide novel sequencing data generated in-house for all technologies. To the best of our knowledge, we provide the first independent evaluation of the ICLR approach. Furthermore, we used the latest truth set release [52] for the evaluation and a cutting-edge comparison approach [53]. This work aims to provide insights regarding the contribution of recent technological advances in human whole genome sequencing, and to serve as a resource for the decision-making in future research projects and technological platforms design and development.

## 2 Methods

### Samples

We used the HG002 human reference for all experiments. For long read sequencing with ONT and PacBio, we used genomic DNA (gDNA) extracted from Lymphoblastoid Cell Line ref. GM24385 (Coriell). The Cell Line was cultured according to the recommendations of Coriell Institute [54], using RPMI 1640 medium (Thermo Fisher), FBS serum (Thermo Fisher) and Penicillin-streptomycin antibiotic (Thermo Fisher). Using KOVA slides (Labelians) we counted 2.0 – 2.5 millions cells per extraction and extracted gDNA with Nanobind CBB kit (PacBio) following the manufacturer’s instructions. For short read sequencing, we used gDNA ref. NA24385 (Coriell). We performed quality control (QC) for all gDNA samples using Nanodrop (Thermo Fisher), Qubit dsDNA High Sensitivity (Thermo Fisher), and TapeStation (Agilent) prior to library preparation.

### Sequencing

We performed Whole Genome Sequencing (WGS) using the following technologies. Each approach was previously implemented at CNRGH in accordance with the manufacturer’s instructions and validated for high throughput production. The conditions described below reflect the protocol validated at CNRGH for production for each approach and are therefore not equivalent among all technologies.

#### Illumina short reads

We prepared the Illumina library from 1.1 µg gDNA input, sheared using LE220plus (Covaris), with 96 microTUBE-50 AFA Fiber Plate (Covaris) and target fragment length 450 bp. For library preparation we used an automated platform Bravo NGS Workstation (Agilent) with TruSeq DNA PCR-Free Library Prep kit (Illumina) and IDT for Illumina TruSeq DNA UD Indexes v2 (Illumina), with clean-up steps using AMPure XP beads (Backman Coulter), all according to the manufacturers’ instructions. We performed QC of the final library using LabChip GX High Sensitivity Reagent kit (Revvity) and KAPA Library Quantification Kit (Roche). We sequenced the library on the NovaSeq X Plus (Illumina) instrument with NovaSeq X Series 25B Reagent kit (300 Cyc) (Illumina) according to the manufacturer’s instructions, using 2×159 bp reads, and 8-plex pooling per lane. We obtained a yield of 131 Gb for this library, with an estimated insert size of 422 bp, a mean quality score of 38.0, and 90.22% of bases Q30+ (demultiplexing and base quality evaluation were performed with bcl-convert 3.9.3.2). The mean depth of coverage without downsampling was 38.07×.

#### ICLR

We prepared the ICLR library from 50 ng gDNA input using Illumina Complete Long Read Prep, Human (Illumina) according to the manufacturer’s instructions. Briefly, the ICLR workflow includes long-range tagmentation (with fragments 5 – 7 kb long), DNA marking by random nucleotide incorporation during long-range amplification (12 PCR cycles), followed by dilution and another long-range amplification (14 PCR cycles), with a clean-up after each step using Illumina Purification Beads (included with the ICLR kit). We performed QC, then we proceeded with tagmentation, indexing PCR (6 PCR cycles) with Illumina Unique Dual Indexes LT (Illumina), clean-up, and final QC. For QC, we used Qubit dsDNA HS assay kit (Thermo Fisher) and 2100 Bioanalyser System (Agilent) with High Sensitivity DNA Kit (Agilent). We sequenced the ICLR library on the NovaSeq 6000 (Illumina) instrument with NovaSeq 6000 S4 Reagent Kit v1.5 (300 cycles) (Illumina) in Xp workflow, with a single library per lane, using 2×151 bp reads and no index reads, according to the manufacturer’s instructions. We obtained a yield of 834 Gb for this library, with an estimated insert size of 582 bp, a mean quality score of 34.49, and 86.43% of bases Q30+. ICLR analysis requires 180 – 200× ICLR data but also 30 – 40× standard Illumina WGS data. We prepared standard Illumina WGS library as described above using TruSeq DNA PCR-Free Library Prep (Illumina) with target fragment length 350 bp. We sequenced the library on the NovaSeq 6000 (Illumina) instrument with NovaSeq 6000 S4 Reagent Kit v1.5 (300 cycles) (Illumina) in standard workflow according to the manufacturer’s instructions, using 2×151 bp reads and 24-plex pooling per flow cell. We obtained a yield of 147 Gb for this library, with an estimated insert size of 373 bp, a mean quality score of 35.86 and 93.9% of bases Q30+ (demultiplexing and base quality evaluation were performed with bcl2fastq 2.20.0.422). We analysed the ICLR together with the standard WGS data using Illumina Complete Long Reads DRAGEN Cloud Analysis, version 1.1.2, in BaseSpace Sequence Hub Professional (Illumina). The mean depth of coverage was 30.87×.

#### PacBio

We prepared the PacBio library from 2.2 µg gDNA input treated with SRE (PacBio) in a volume of 77 µL, sheared using Megaruptor 3 (Diagenode) at speed 29 in a volume of 100 µL, using SMRTbell Prep kit 3.0 and Revio Polymerase kit (PacBio), with clean-up steps using AMPure PB beads (PacBio), all according to the manufacturers’ instructions. We performed QC using Qubit dsDNA HS assay kit (Thermo Fisher) and Femto Pulse gDNA 165kb Analysis Kit (Agilent). We sequenced the library on the Revio instrument (PacBio) using one Revio SMRT Cell (PacBio) per individual, with the options of adaptive loading and movie time 30h selected in SMRT Link v13.1. We obtained a yield of 118 Gb HiFi for this library, with a median read length of 18 275 bp, a mean quality score of 38.37, and 92.78% of bases Q30+. The mean depth of coverage without downsampling was 39.69×.

#### ONT

We prepared the ONT library from 1.0 µg gDNA input (with no short read eliminator treatment), sheared using Megaruptor 3 (Diagenode) at speed 33 in a volume of 100 µL, using Ligation Sequencing kit V14 (ONT) and NEBNext Companion Module for ONT Ligation Sequencing (NEB), with clean-up steps using AMPure XP Beads (included with the ONT Ligation kit), all according to the manufacturers’ instructions. We performed QC using Qubit dsDNA HS assay kit (Thermo Fisher). We sequenced the ONT library on the PromethION 24 instrument (ONT) using PromethION flow cell R10.4.1 (ONT), with a 72h sequencing run, with one flow cell wash and reload. We performed onboard basecalling in FAST mode in MinKNOW Core version 5.8.6 and MinKNOW PromethION release version 23.11.7. After the run, we performed SUP basecalling in Dorado version 0.3.4 (model dna r10.4.1 e8.2 400bps 5khz sup.cfg). We obtained a yield of 113 Gb for this library, with a median pass read length of 15 842 bp, and a mean quality score of 29.05. The mean depth of coverage without downsampling was 32.63×.

### Variant discovery pipeline

The bioinformatics analyses are summarized in Fig. 1 and described below for Illumina short reads, PacBio, and ONT. We used widely adopted, reference bioinformatics tools for each technology. For ICLR data, all steps from read mapping to variant calling were performed using Illumina Complete Long Reads DRAGEN Cloud Analysis [v1.1.2] [55].

**Fig. 1.**
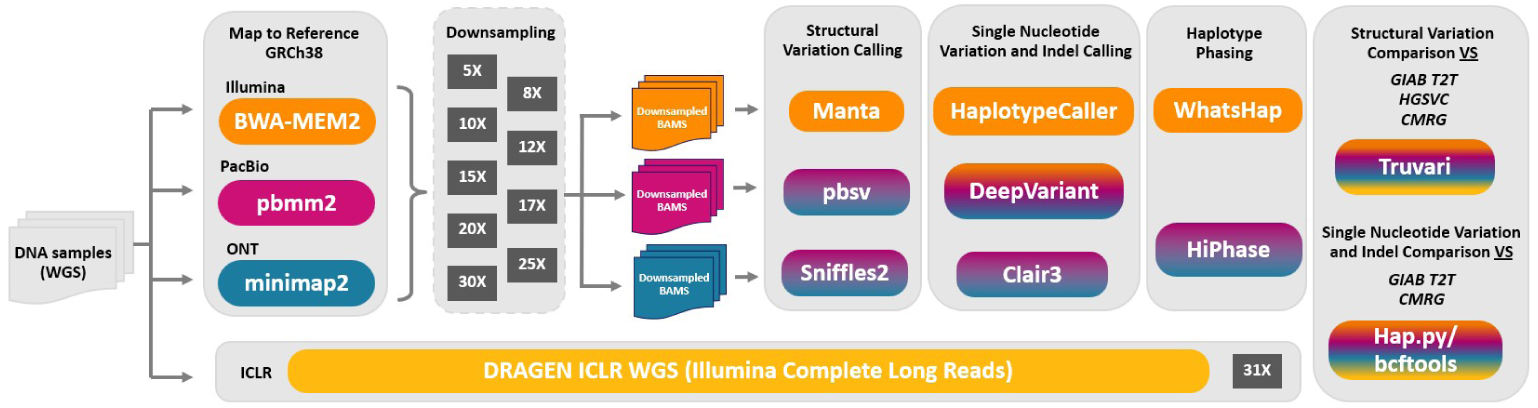
Bioinformatic pipeline for whole genome analysis. The different steps include mapping, downsampling, structural variation calling, small variant calling, phasing, and comparison to the following truth sets: Genome in a Bottle Telomere-to-Telomere (GIAB T2T), Challenging Medically Relevant Genes (CMRG), and The Human Genome Structural Variation Consortium (HGSVC).

### Mapping

We mapped the short and long read sequencing data to the GRCh38 reference using different alignment tools optimized for each technology (Supplementary Template 1). Illumina short reads, known for their high accuracy but short read length, were mapped using BWA-MEM2 [v2.2.1], a fast and memory-efficient aligner designed for short read mapping [21], with the option -Y that allows soft clipping for supplementary alignments. PacBio HiFi reads, which are highly accurate long reads, were aligned using pbmm2 [v1.13.1] [56], a PacBio-optimized wrapper for minimap2 [57, 58] with the option --preset HIFI to improve alignment accuracy enhancing structural variant detection and phasing. ONT long reads, which can span megabases but have higher error rates, were mapped using minimap2 [v2.26], a versatile aligner designed for higherror long reads. We used the parameter -x map-ont, which is a preset for aligning ONT reads to reference genomes. This mapping step ensures that sequencing reads from different platforms were accurately aligned to the reference genome for downstream analysis. We performed post-alignment quality control using samtools [v1.15.1] [23, 59], seqkit [v2.8.1] [60] and bedtools [v2.30.0] [61].

### Downsampling

The mapped reads were downsampled to different coverage depths using the subsample option in samtools [v1.15.1] (Supplementary Template 2). The goal was to create subsets of the original data at various coverage depths, specifically 5×, 8×, 10×, 12×, 15×, 17×, 20×, 25×, and 30× for PacBio, ONT, and Illumina. This approach allows the evaluation of sequencing performance and analyses at different coverage levels. It provides insights into the minimum coverage required for variant calling and haplotype phasing as well as comparing the effectiveness of the different sequencing technologies at equivalent depths. We selected the same downsampling points as [50] to allow direct comparisons with this study. ICLR could not be downsampled due to the limitations of ICLR data processing in DRAGEN.

### Structural variant calling

We performed SV calling from downsampled data to assess the sensitivity and the accuracy of different SV callers at equal depth of coverage. This approach helps determine the minimum coverage required for reliable SV identification. Three SV callers were used in this analysis: Manta [v1.6.0] [36], Sniffles2 [v2.2] [62] and pbsv [v2.9.0] [48] (Supplementary Template 3). Manta is designed for Illumina short read data. The used --callRegions=*{*call regions*}* option restricts variant calling to specific genomic regions, improving efficiency and reducing unnecessary computations. For long read sequencing, Sniffles2 and pbsv were used for both PacBio HiFi and ONT reads. Sniffles2 runs in a single-step process relying on realignment-based methods and statistical filtering to improve SV detection across different read types. Pbsv uses a k-mer-based approach optimized for PacBio HiFi reads (--hifi flag for PacBio). It follows a twostep process: (1) pbsv discover generates an intermediate SV signature file from the BAM file and (2) pbsv call uses the reference genome and SV signature file to call SVs. Manta is optimized for short reads and has a simpler configuration, whereas Sniffles2 and pbsv require additional settings for long-read-specific features such as tandem repeats annotations to improve calling in repetitive regions, alignment lengths, minimum support reads, and MAPQ scores.

### Small variant calling

We conducted SNV and indel calling from downsampled data using HaplotypeCaller program [63] in GATK [v4.6.0.0] for short read sequencing data [24], Clair3 [v1.0.5] for long read sequencing data [27] and DeepVariant [v1.6.1] for all the three technologies [26] (Supplementary Template 4).

To perform small variant calling from Illumina short reads, GATK provides a two-step process. First, CalibrateDragstrModel is used to estimate parameters for the DRAGstr model, which helps in analyzing short tandem repeats (STRs). Then, HaplotypeCaller is executed utilizing the calibrated DRAGstr model for small variant calling. This step performs local *de novo* assembly of haplotypes in active regions, enabling simultaneous calling of SNVs and indels.

Clair3 is a high-performance small variant caller designed for long read sequencing data. It combines the strengths of pileup and full-alignment methods to achieve superior accuracy and speed in small variant calling. The Clair3 command is relatively similar for both technologies except for two settings: --platform=hifi*|*ont and --model path. The platform-specific models ensure optimized performance for the unique characteristics of each technology. For PacBio HiFi data, we used the specific “hifi revio” model, which is designed for the latest PacBio Revio sequencer. For ONT, we used the “r1041 e82 400bps sup v420” model, which is tailored for ONT sequencing. Both commands enable phasing, which can enhance SNV and indel calling accuracy.

DeepVariant is a deep learning-based variant caller that can be used with different sequencing technologies. Each command is tailored to the specific characteristics of the sequencing platform. For Illumina data, DeepVariant uses the ‘WGS’ model type and includes additional parameters for haploid contigs and pseudo-autosomal regions on sex chromosomes. The PacBio command utilizes the ‘PACBIO’ model type, specifically designed for long read data from this platform. For ONT, the command employs the ‘ONT R104’ model, which is optimized for the R10.4 chemistry used in ONT sequencing. The Illumina command includes extra arguments for read processing, while the PacBio and ONT commands are more streamlined. These differences highlight DeepVariant versatility in handling diverse data types for accurate small variant calling.

### Phasing

Haplotype phasing is the process of determining which genetic variants are inherited together on the same chromosome, crucial for understanding genetic inheritance patterns and identifying disease-causing mutations. In this study, we performed phasing on downsampled sequencing data from Illumina, PacBio, and ONT (Supplementary Template 5). For Illumina short read data and ICLR, small variants were phased using WhatsHap [v2.2] [64]. While it was originally designed to fully exploit long reads from PacBio and ONT, it also performs effectively with high-accuracy Illumina short reads. Three programs were applied (whatshap phase, whatshap stats and whatshap haplotag). This software uses a haplotype reconstruction model based on the minimum error correction (MEC) principle, which aims to assign reads to haplotypes while minimizing inconsistencies. For PacBio and ONT long read sequencing data, small and structural variants were phased jointly using HiPhase [v1.4.0] [65], which is designed to leverage the long-range information provided by these technologies. HiPhase employs a dual-mode allele assignment approach and a novel application of the A* search algorithm to solve the phasing problem efficiently. It demonstrates superior performance by producing longer and more accurate phase blocks than many existing tools. We calculated the breadth of coverage of phased genome as the ratio of Total block size to the reference genome size (2 934 876 545 bp).

### Assessment of structural variant calling

To assess the accuracy of the called structural variants, we compared the results against high-confidence truth sets, including the Genome in a Bottle Telomere-to- Telomer high-confidence set (GIAB T2T) [66, 67], Challenging Medically Relevant Genes (CMRG) [68, 69], and The Human Genome Structural Variation Consortium (HGSVC) [70, 71]. The three truth sets were filtered with bcftools to keep only insertions and deletions larger than 50 bp that passed all quality filters (Supplementary Table S1).

The benchmarking process was carried out using Truvari [v4.3.1] [53], a toolset designed for structural variant comparison. Specifically, Truvari’s components – bench, refine, and ga4gh – were employed to evaluate the precision, recall and F1 score of the SV calls against the two SV truth sets GIAB T2T and HGSVC. Only the Truvari bench method was performed for CMRG. This truth set is designed for direct benchmarking, meaning the variants are already preprocessed and carefully curated. Our evaluation is based on the precision, recall, and F1 scores as defined in Supplementary Section 3.

The Truvari bench command evaluates the accuracy of VCF files by comparing it to a truth set using a reference genome, in our case GRCh38. The benchmarking was restricted to specific genomic regions, available from [72] for GIAB T2T and HGSVC, and from [73] for CMRG.

Following the GIAB T2T reference evaluation parameters with several adjustments, only high-quality variants were considered (--passonly), variants smaller than 50 bp were ignored with relaxed matching criteria (variants are considered matching if they are within a maximum distance of 2000 bp, and all variants within a chunksize of a maximum distance of 5000 bp are grouped for comparisons). Duplications were treated as insertions, and in case of multiple matches, the variant with the highest allele count (--pick ac) was selected (Supplementary Template 6).

The Truvari refine method is a valuable tool for improving the accuracy of SV benchmarking by harmonizing phased variants with MAFFT local alignment [74]. Building upon initial benchmarking results, it restricts the analysis to candidate regions, enabling a more precise and comprehensive comparison of SVs. This refinement process, outlined in the command line in Supplementary Template 7, relies on phased SV information derived from HiPhase, specifically using long read sequencing data. Long reads are particularly advantageous for accurately phasing large SVs due to their ability to span extended genomic regions, thereby improving the reliability of SV calls in complex genomic contexts.

Truvari ga4gh is an important tool for ensuring that SV benchmarking results adhere to the standards set by GA4GH (Global Alliance for Genomics & Health) [75], which aim to enhance consistency, interoperability, and reproducibility in genomics research [76]. The inclusion of the --with-refine option enables Truvari to replace variants with refined versions in regions where harmonization succeeds, while keeping the original variants unchanged in regions where harmonization is not possible. Precision, recall and F1 score are thus recomputed on the refined set of comparable variants. Command line is shown in Supplementary Template 7.

### Assessment of small variant calling

To measure the robustness of small variants detection, we employed hap.py [v0.3.15] [77], a widely used tool designed for benchmarking SNV and indel calls against high-confidence gold standard truth datasets. Unlike simple row-by-row VCF record comparisons, hap.py performs a haplotype-level comparison of diploid genotypes, providing a more accurate and biologically meaningful assessment of small variant calling performance.

In this study, we compared our small variant call set against GIAB T2T and CMRG. We performed the comparison using the RTG vcfeval engine [v3.12.1] [78], following the GA4GH best practices for variant evaluation. The analysis was restricted to high-confidence genomic regions defined in the GIAB T2T benchmark BED file, using the GRCh38 reference genome. Additionally, a stratification file was incorporated to analyze the performance across different variant categories and the sample was specified as male to account for sex chromosome differences (Supplementary Template 8). This benchmarking approach ensures a robust evaluation of small variant calling performance, providing insights into precision, recall, and F1 score. An additional bcftools [v1.15.1] [59] step was added to extract statistics from the obtained VCF files.

### Phasing assessment

Phasing accuracy was evaluated using *WhatsHap compare* by comparing phased variant calls for HG002 against the GIAB T2T truth set (Supplementary Template 9). This tool aligns inferred haplotypes to the truth set and reports, for each chromosome, the number of switch errors and the total number of phase-informative adjacent variant pairs assessed. The switch error rate (SER), defined as the ratio of switch errors to assessed variant pairs, provides a measure of local phasing accuracy.

SERs were computed for Illumina, PacBio, and ONT (30× coverage) and ICLR (30.87× coverage) by comparing phased variant calls against the GIAB T2T truth set. Using bcftools, all phased VCF files were restricted to autosomal chromosomes, compressed, indexed, and further filtered to retain only biallelic variants to ensure compatibility with switch error computation. The global SER was computed as the total number of switch errors divided by the number of consecutive heterozygous variant pairs jointly phased in both query and reference datasets.

## 3 Results

### Quality metrics overview

To characterize the sequencing output generated by the different technologies, we evaluated the read length, sequence accuracy and the coverage organization. The insert size for Illumina (Fig. 2A) displays a tight distribution centered around a median of 422 bp, corresponding to the fragmentation target size at 450 bp. ICLR technology yields longer reads than standard Illumina short reads, with a median length of 4 kb in a bimodal distribution that is the expected shape for ICLR data. The observed median length is somewhat below the fragmentation target size of 5 – 7 kb. Both long read technologies share close median read lengths, 16 kb for ONT and 18 kb for PacBio, reflecting their slightly different fragmentation setting. For both technologies, 90% of all reads are below 27 kb. While PacBio generates a tighter distribution between 517 bp and 62 kb, ONT sequencing provides a wider length range from 87 bp up to nearly 962 kb, with an important tail of short fragments due to the absence of short fragment elimination step during library preparation.

**Fig. 2.**
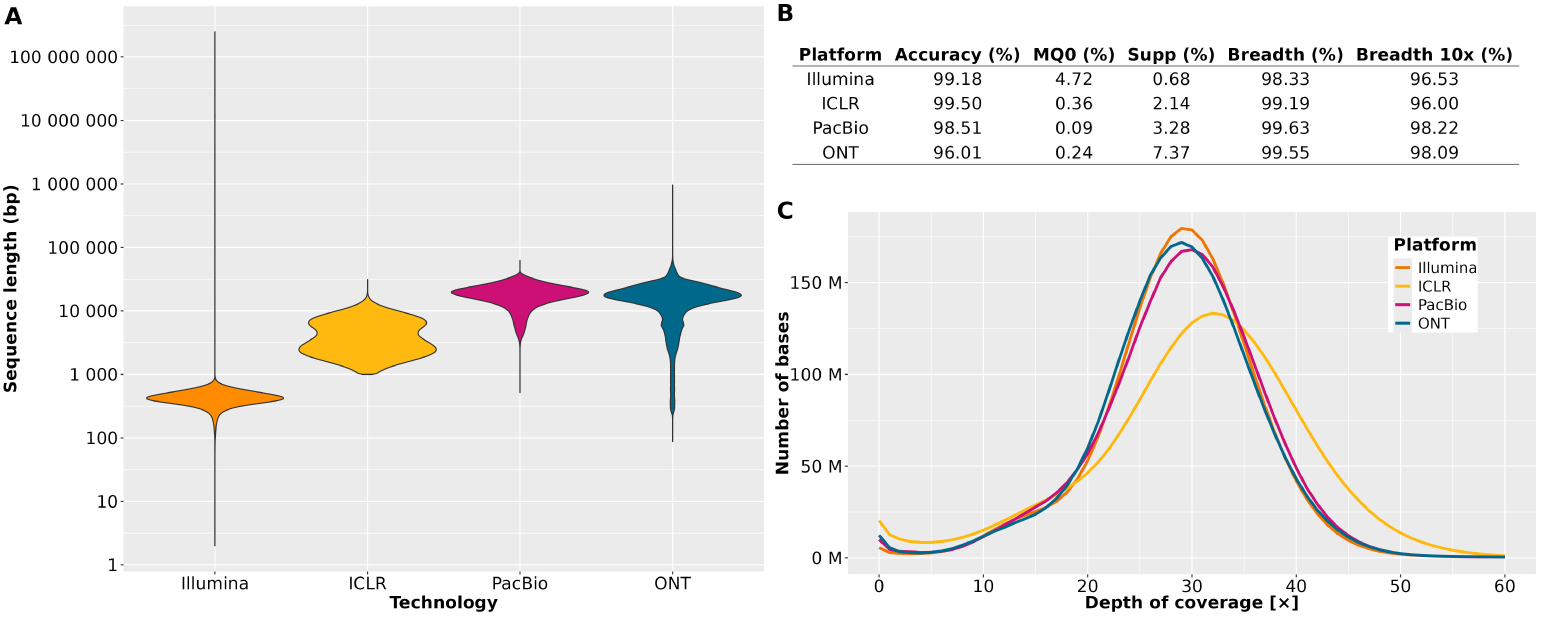
Quality control overview after mapping to GRCh38 reference at 30× mean depth of coverage. Illumina - Illumina short reads, ICLR - Illumina Complete Long Reads, PacBio - Pacific Biosciences, and ONT - Oxford Nanopore Technologies. **(A)** Violin plots of the read length distribution (insert size in case of Illumina short reads). **(B)** Quality control metrics: MQ0 - the proportion of reads with mapping quality at 0, Supp - the proportion of reads tagged as supplementary, Breadth - the breadth of the genome covered with at least one read, Breadth10x - the breadth of the genome covered with at least 10 reads. **(C)** Coverage evenness. Please note that for ICLR the mean coverage is at 30.87× due to technical constraints of DRAGEN analysis, while it is 30× for all other technologies.

Sequence accuracy of mapped reads varies among technologies (Fig. 2B). Illumina short reads and ICLR achieve the highest level of accuracy, both above 99%. PacBio provides sequence accuracy close to short reads (98.51%). Despite constant evolution of the technology and bioinformatics, particularly signal processing and post-analysis, ONT still displays the lowest accuracy (96.01%).

We evaluated the coverage organization after downsampling all datasets to an equal mean depth of coverage (30×), except for ICLR, which could not be downsampled freely in DRAGEN and was therefore evaluated at the original mean depth of 30.87×. The breadth of coverage shows the advantage of long read technologies (Fig. 2B): PacBio and ONT provide the highest breadth of coverage (99.63% and 99.55%, respectively), closely followed by ICLR (99.19%), compared to Illumina short reads (98.33%). Long reads provide coverage in complex regions that are inaccessible with short reads, as illustrated by the difference of MQ0 between Illumina short reads (4.72%) and other technologies: PacBio yielded the lowest MQ0 (0.09%), ONT yielded the second lowest MQ0 (0.24%, which is however associated to the highest proportion of supplementary alignments, presumably due to alternative alignments of the longest reads on the current reference), followed by ICLR with MQ0 at 0.36%. Coverage evenness is similar among all technologies, sharing similar bell-shaped depth of coverage profiles as expected for whole genome sequencing, which is centered around the mean depth of coverage value (Fig. 2C). Overall, long read sequencing yields broader coverage of the genome, therefore providing better material for downstream analysis.

### Structural variant detection

Structural variation includes different types of genome modifications of wide range of sizes and complexities. Short read sequencing has limited power to characterize large rearrangements as well as events in complex regions, such as repetitive regions or genomic regions highly divergent from the reference genome. In this context, SV detection is highly improved by the use of long read sequencing technologies (Fig. 3A): ICLR, PacBio, and ONT provide almost a three-fold increase in the number of detected SVs (24 571, 23 492, and 23 405 SVs, respectively) compared to Illumina short reads (8 198 SVs). Long read technologies are particularly better adapted than short reads for the detection and characterization of insertions, since short reads cannot span the whole length of large inserted or modified sequences. Inversions, although sparsely represented in the human genome, are likewise better detected with long read approaches. The number of uncharacterized breakend events detected by each technology indicates the level of ambiguity in the SV type classification and the definition of their boundaries. Currently, long read technologies provide the lowest proportion of breakend events, demonstrating better SV definition and annotation (Fig. 3A). In a detailed comparison among long reads, the SV profiles are very similar between ONT and PacBio, differing slightly by a higher number of duplications detected by PacBio. We attribute this slight difference to bioinformatics processing specificities: Fig. 3A shows the SV profiles of PacBio with pbsv caller recommended by PacBio while Supplementary Fig. S2 shows an alternative with Sniffles2. In contrast, the SV profile of ICLR is different, with more observed breakend events and deletions, and fewer insertions.

**Fig. 3.**
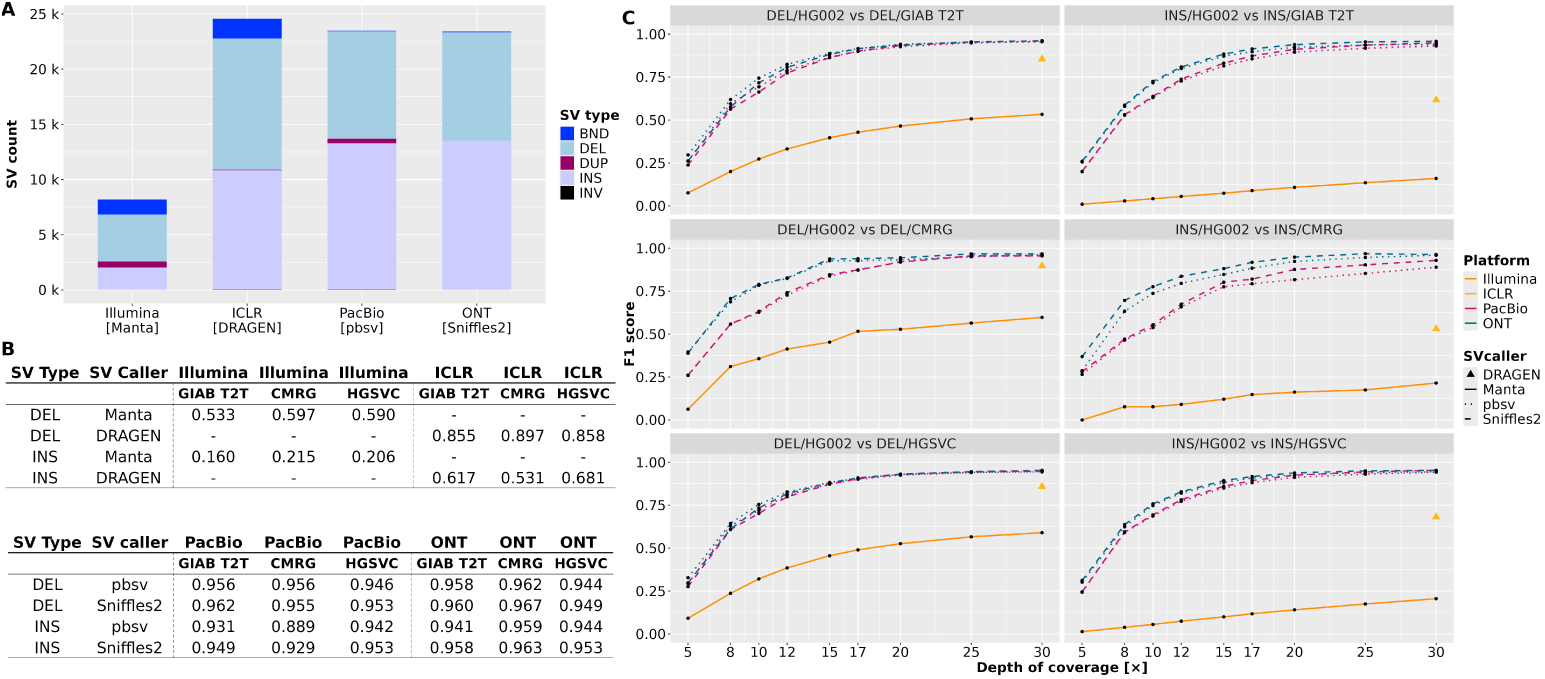
Results of structural variant (SV) profiling across all sequencing technologies. Illumina - Illumina short reads, ICLR - Illumina Complete Long Reads, PacBio - Pacific Biosciences, and ONT - Oxford Nanopore Technologies. **(A)** The counts of different SV types detected at 30×: BND - breakend events, DEL - deletions, DUP - duplications, INS - insertions, and INV - inversions. **(B)** F1 scores for DEL and INS at 30× compared to the following truth sets: Genome in a Bottle Telomere-to-Telomere (GIAB T2T), Challenging Medically Relevant Genes (CMRG), and The Human Genome Structural Variation Consortium (HGSVC). **(C)** The decline of SV calling performance (F1 score) with decreasing mean depth of coverage for DEL and INS for all three SV truth sets.

To evaluate the accuracy of SV profiling, we compared the detected SVs to three different truth sets: GIAB T2T, CMRG and HGSVC, as described in Supplementary Data (Supplementary Table S1 and Fig. S1). Illumina short read sequencing yields F1 score of 0.533 for deletions and only 0.160 for insertions when comparing to the GIAB T2T truth set (Fig. 3B), mostly due to the large number of false negatives (6.0k deletions and 14.2k insertions). ICLR sequencing technology shows intermediate performance with important improvement over Illumina short reads, with F1 score of 0.855 and 0.617 for deletions and insertions, respectively. Despite the equivalent number of SVs detected by ICLR, ONT, and PacBio, the F1 score for ICLR is below that of other long read technologies, both for deletions and insertions. Specifically for insertions, the difference is substantial due to the large number of false negatives (5.6k). PacBio and ONT show good performance for both insertions and deletions, with F1 scores of 0.949 – 0.962 when comparing to the GIAB T2T truth set (the number of false negative insertion calls with Sniffles2 is 1.1k for PacBio and 0.9k for ONT). The results are consistent for both SV callers used in this study, with Sniffles2 providing slightly higher F1 scores than pbsv, due to the higher number of false positives detected with pbsv. When comparing to the CMRG truth set, which contains a higher density of complex regions, ONT achieves slightly better results than PacBio, with insertions F1 score of 0.963 for ONT and 0.929 for PacBio. Supplementary Table S2 shows the number of SV calls shared between short and long reads.

We evaluated the resilience of SV profiling against a progressive decrease in depth of coverage. We downsampled Illumina, PacBio and ONT datasets to a mean depth of coverage ranging from 5× to 30×. ICLR could not be downsampled due to the limitations of ICLR data processing in DRAGEN. All three technologies share a similar SV calling profile with relatively stable performance down to 20×, and progressive decline of F1 score below 15× mean depth of coverage. In the complex regions of the CMRG truth set, SV profiling with ONT is more resilient than PacBio to decreasing depth of coverage, with a faster decline of the PacBio F1 score below 20× for both insertions and deletions. In other words, in the context of this use case, ONT provided globally a better SV profiling for lower depths of coverage.

Next, we evaluated the impact of SV size on the capacity of detection at different depths of coverage. We divided all deletions and insertions by size into four classes (SV counts are indicated in Supplementary Fig. S1). The size classes are arbitrary but they reflect different types of SV annotations (small SVs, SINE, LINE, and large SVs). Fig. 4 shows the F1 scores compared to the GIAB T2T truth set. For deletions, long read technologies generally perform better than short reads. PacBio and ONT show similar profiles regardless of the deletion size, with F1 score around 0.961 except for the longest deletions where F1 score drops slightly to around 0.820. This pattern is mirrored by ICLR, with stable F1 score around 0.872 in all size classes except a slight drop to 0.727 for the longest deletions. In contrast, Illumina short reads show a different pattern: detection improves with increasing deletion size, starting from an F1 score of 0.461 for the shortest deletions and reaching 0.769 for deletions above 1 000 bp. This observation can be explained by the fact that in case of short reads, smaller deletions are mostly confronted with mapping issues, while larger deletions are better supported by the abrupt change of the depth of coverage at the deletion boundary, with larger deletions leveraging a better detection. Compared to the CMRG and HGSVC truth sets (Supplementary Fig. S3 and Fig. S4), F1 scores show similar trends to that described here for GIAB T2T.

**Fig. 4.**
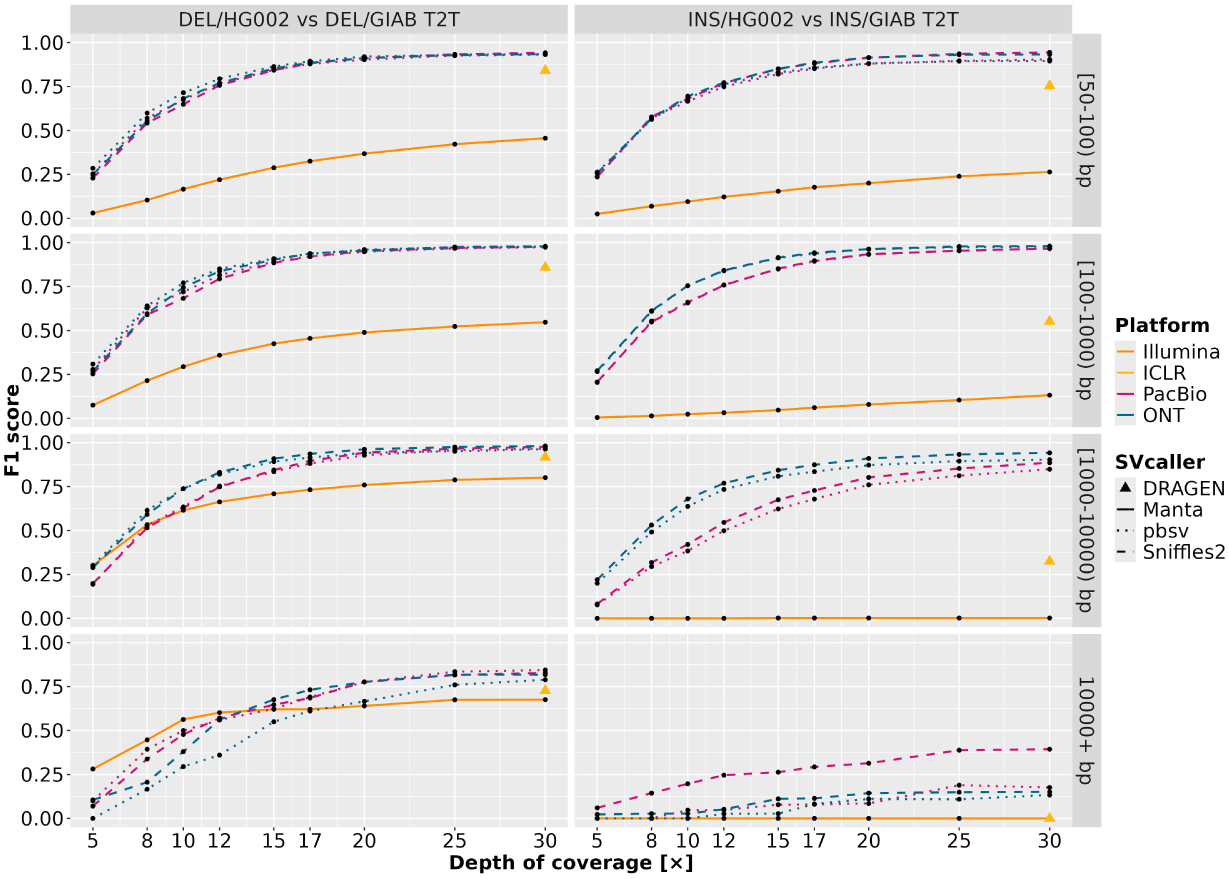
Comparison of SV calling performance by coverage and variant size classes. F1 scores were calculated by progressively downsampling mean depth of coverage for deletions (DEL) and insertions (INS) across four structural variant size categories. Results were benchmarked against the Genome in a Bottle Telomere-to-Telomere (GIAB T2T) truth set over all sequencing technologies: Illumina - Illumina short reads, ICLR - Illumina Complete Long Reads, PacBio - Pacific Biosciences, and ONT - Oxford Nanopore Technologies.

For insertions, long read technologies perform far better than short reads, regardless of the insertion size (Fig. 4). As described above, Illumina short reads generally provide poor description of insertions, which is aggravated for larger insertions, starting at F1 score of 0.253 for the smallest insertions and dropping to F1 score of 0 for insertions above 1 000 bp. ICLR also shows a diminution of the F1 score with increasing insertion size, with F1 score of 0.754 for the shortest range and dropping to 0 for insertions above 10 000 bp. ONT and PacBio performance is also impacted by the insertion size, starting from F1 score around 0.946 for the insertions below 1 000 bp, decreasing slightly for insertions between 1 000 and 10 000 bp long in case of PacBio, and dropping dramatically for insertions above 10 000 bp long in case of both PacBio and ONT to around 0.153, except for PacBio insertions called with Sniffles2, which performs slightly better at 0.394. This observation highlights the fact that for the largest insertions, detection is highly dependent on the bioinformatics processing applied. Even with the advantage of long read technologies, many large insertions remain undetected with mapping-based approach: an alternative assembly-based approach to SV calling may be better adapted for very large insertions.

On the whole, this evaluation shows that long reads sequencing represented by ONT and PacBio provide a better SV profiling than ICLR and Illumina short reads, in terms of accuracy, completeness, and resilience to reduced depth of coverage. While globally the results generated by SV callers for long read data share similar profiles, the choice of bioinformatics tools may introduce specific variations for certain SV profiles, especially for larger or complex events.

### Small variants detection

We examined the quality of small variant profiling (SNVs and indels *<*50 bp long) to assess the performance of each technology compared to two reference datasets: the GIAB T2T and the CMRG truth sets.

Fig. 5A shows the total number of SNVs, ranging from 3.9M detected by Illumina short reads, through ICLR and ONT each detecting approximately 4.1M SNVs, to 4.3M detected by PacBio. Compared to the GIAB T2T truth set (Fig. 5B), ICLR provides the lowest total number of false negatives and false positives (24k and 8k, respectively), which is likely due to the large amount of data used (*>*200×) to generate the consensus sequence during the specific processing applied in DRAGEN. ICLR is closely followed by PacBio, providing the second-lowest total number of false negatives and false positives (20k and 18k, respectively), leveraging the advantage of long reads of better mappability in complex regions, as well as high sequence accuracy of PacBio technology. Both ICLR and PacBio yield the highest F1 scores for both truth sets (0.990 - 0.996) (Fig. 5C). SNV profiling from Illumina short reads shows the highest number of false negatives (55k), likely due to its limited mappability in complex regions, while the number of false positives remains low (16k), likely due to high sequence accuracy. The F1 score of Illumina short reads shows a difference between GIAB T2T (0.990) and CMRG (0.974), indicating difficulties in SNV calling in complex regions. Despite the advantage of better mappability in complex regions over short reads, ONT also shows many false negative SNV calls (50k) and the highest number of false positives (32k), likely due to inferior accuracy, or possibly due to the variation of read size distribution locally available for SNV calling in complex regions.

**Fig. 5.**
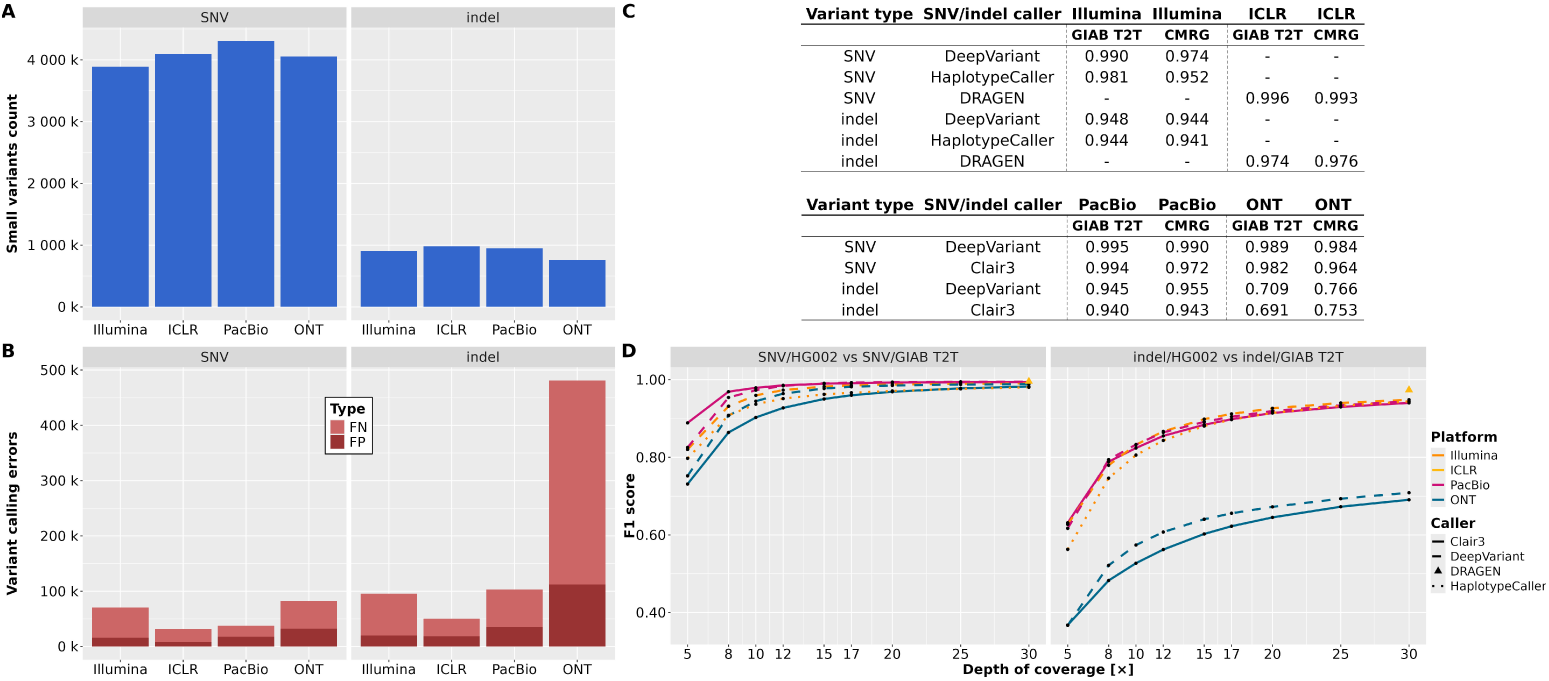
Results of small variant profiling, covering single nucleotide variants (SNVs) and indels (*<*50 bp). Illumina - Illumina short reads, ICLR - Illumina Complete Long Reads, PacBio - Pacific Biosciences, and ONT - Oxford Nanopore Technologies. **(A)** The counts of SNVs and indels detected at 30×. **(B)** The counts of variant calling errors for SNVs and indels at 30× compared to Genome in a Bottle Telomere-to-Telomere (GIAB T2T) truth set: false negative (FN) and false positive (FP) calls. **(C)** F1 scores for SNVs and indels at 30× compared to the following truth sets: Genome in a Bottle Telomere-to-Telomere (GIAB T2T) and Challenging Medically Relevant Genes (CMRG). **(D)** The decline of small variant calling performance (F1 score) with decreasing mean depth of coverage for the GIAB T2T truth set.

Regarding indels, we observe a striking difference between ONT and all other technologies, with ONT yielding the highest number of false negative and false positive indels (369k and 112k, respectively, Fig. 5B). Despite a dynamic evolution of com- putational models used for ONT basecalling and indel calling, our observations seem to indicate that ONT technology remains exposed to difficulties in complex regions, resulting in substantial limitations in indel profiling. PacBio and Illumina short reads share similar profiles, both performing substantially better than ONT with approximately 100k total count of false negative and false positive indel calls. PacBio yields fewer false negatives than Illumina, likely due to the better mappability of long reads in complex regions, while Illumina yields fewer false positives than PacBio, likely related to better sequence accuracy in correctly mapped regions. ICLR shows the fewest false negative and false positive indel calls (50k in total), again likely due to the large amount of sequencing data used to generate the dataset.

We evaluated the impact of different bioinformatics tools on the performance of SNV and indel calling, compared to GIAB T2T and CMRG truth sets (Fig. 5C). In general, the tendencies described above were comparable, regardless of the bioinformatics tools used, and for both truth sets. DeepVariant provided systematically the best results for SNV and indel detection for all sequencing technologies.

Next, we evaluated the resilience of small variant detection against a progressive decrease in depth of coverage. Both the GIAB T2T and CMRG truth sets comparisons yield similar results (Fig. 5D and Supplementary Fig. S5): SNV calling shows very close results for all technologies, and fairly resilient to downsampling, with the F1 score starting to drop below 12× mean depth of coverage. DeepVariant performs better than both Clair3 and HaplotypeCaller in SNV calling from complex regions of the CMRG truth set. Indel profiling is less resilient to downsampling than SNV profiling. Regardless of the sequencing technology and bioinformatics processing, the F1 scores for indels start to drop for any coverage below 20×.

In summary, all technologies evaluated here demonstrated the capacity to provide accurate SNV profiles for depth of coverage ranging from 12× to 30×. Illumina provides the highest accuracy, while PacBio provides the best breadth of coverage and subsequently a better completeness of variant detection. For indels, Illumina short reads, ICLR, and PacBio all provide accurate indel profiles between 20× and 30× coverage but the performance drops at lower coverage. In contrast to other technologies, ONT still needs to improve before it can be considered as an appropriate solution for indel detection, regardless of coverage.

### Phasing

The phasing of genomic variants is a relevant input for genome variant profiling, notably supporting a better understanding and detection of compound heterozygosity and genetic burden analysis. Haplotype phasing information has been underused because of the limitations of short read technologies. However, with the advent of long read technologies, phasing can provide important insights into genetic analyses and disease modelization. The performance in phasing shows major differences among the evaluated technologies (Table 1). In general, long read technologies provide better phasing and higher assembly contiguity than short reads. With long reads, we observe a considerable increase in the phased block NG50 length, starting from null for Illumina short reads, with modest performance of ICLR at 95 kb, reaching 535 kb for ONT and 720 kb for PacBio. This NG50 value suggests that due to its inherent limitations, Illumina short read technology did not produce a sufficient number of high-quality contigs or scaffolds. We observe a substantial diminution in the number of phased blocks with long reads, starting from 506k for Illumina short reads, with intermediate performance of ICLR with 33.8k phased blocks, reaching 10.8k and 7.7k for ONT and PacBio, respectively. Long reads result in lower phasing sparsity, with the breadth of coverage of phased genome over 80% for both ONT and PacBio.

**Table 1.** Summary of haplotype phasing performance at 30× mean depth of coverage. Illumina - Illumina short reads, ICLR - Illumina Complete Long Reads, PacBio - Pacific Biosciences, and ONT - Oxford Nanopore Technologies.

|  | Number of blocks | Max block size (bp) | Total block size (bp) | Block NG50 (bp) | Switch error rate (%) |
| --- | --- | --- | --- | --- | --- |
| Illumina | 505 512 | 95 216 | 339 130 602 | 0 | 1.4448 |
| ICLR | 33 854 | 1 786 742 | 1 939 829 830 | 95 381 | 0.6832 |
| PacBio | 7 659 | 18 331 778 | 2 463 070 368 | 719 859 | 0.0619 |
| ONT | 10 775 | 30 127 848 | 2 419 195 617 | 535 142 | 0.1058 |

In addition to genome contiguity, phasing accuracy was assessed using the switch error rate (SER), a standard metric for quantifying local phasing errors. At 30× mean depth of coverage, long read technologies consistently exhibited lower SERs than short read data, indicating superior local phasing accuracy. Among the evaluated technologies, PacBio achieved the lowest SER, followed by ONT. ICLR showed an intermediate SER, higher than long read technologies but lower than Illumina short read data.

While ONT and PacBio share similar phasing metrics, differences appear in the distribution of block size. ONT longer reads leverage the building of larger phased blocks illustrated by the maximal block size of 30 Mb for ONT against 18 Mb for PacBio. Nonetheless, with more centered read length distribution and higher sequence accuracy, PacBio enables globally a better phasing with a longer phased block NG50 and fewer phased blocks. The comparatively poor performance of ICLR in phasing is likely due to the initial DNA fragmentation size of 4 kb, which is four-fold shorter than ONT and PacBio, thus limiting the capacity to link reads through complex regions and provide the most comprehensive phasing.

We evaluated the resilience of phasing across progressively decreasing sequencing depths using the downsampling approach described above. This analysis was restricted to PacBio and ONT for the following reasons. Illumina short reads were excluded due to a null NG50 value at 30× coverage. ICLR dataset could not be downsampled because of DRAGEN Illumina pipeline limitations previously mentioned. Phasing performance decreased proportionally with sequencing depths for all technologies, for example at 20× mean depth of coverage, the NG50 length decreases to 456 kb for ONT and 599 kb for PacBio (Fig. 6).

**Fig. 6.**
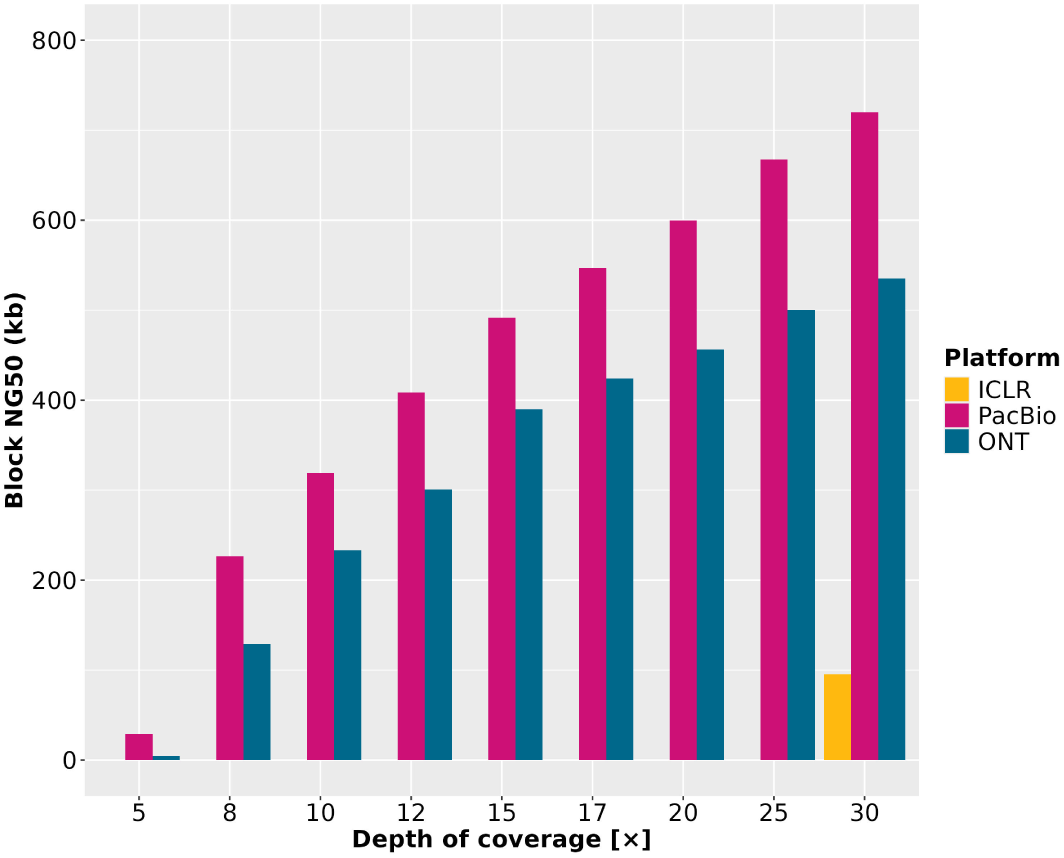
The decline of the block NG50 length with decreasing mean depth of coverage using HiPhase for Pacific Biosciences (PacBio) and Oxford Nanopore Technologies (ONT), and WhatsHap for Illumina Complete Long Reads (ICLR).

These results highlight the importance of long read sequencing for accurately resolving structural genome organization and achieving high-quality haplotype phasing.

## 4 Discussion

This work investigates the performance of different sequencing technologies in human whole genome sequencing and variant profiling. We benchmarked Illumina short reads, long read technologies represented by ONT and PacBio, and Illumina’s reconstructed long reads approach ICLR.

In general, all technologies performed well in SNV profiling, with ICLR and PacBio yielding the highest F1 scores, followed closely by Illumina short reads and ONT. Illumina short reads appear to have the advantage over long reads in precision, with fewer false positive calls owing to the high sequence accuracy. In contrast, long reads appear to have the advantage in recall, because Illumina yields more false negative calls in regions of low mappability. This observation is compatible with previous findings [4]. For SNV profiling in CMRG regions, ICLR, PacBio and ONT all outperformed Illumina short reads, confirming the advantage of long reads, which provide better access to complex regions of the human genome such as segmental duplications and low-mappability regions [68]. ICLR, Illumina short reads and PacBio yielded reliable results in indel profiling. In contrast, in spite of using the ONT latest R10 chemistry with improved accuracy, we observed that indel calling with ONT approach remains inferior to other technologies. This observation thus remains unchanged, compared to previous studies based on less accurate R9 chemistry, which compared ONT to Illumina short reads [4] and PacBio [50].

Long read technologies provided a three-fold increase in the number of detected SVs compared to Illumina short reads. PacBio and ONT both provided the most accurate SV profiling, as expected [10, 12, 16, 44, 70]. In SV profiling, ICLR showed an intermediate performance between short and long reads, which fell short of PacBio and ONT even for the shortest class of SVs. In summary, the current capacity of short reads to correctly identify SVs is limited. In this study, Illumina short reads missed ~13 000 true SVs compared to PacBio and ONT, representing an approximate cumulative size of 4.5 Mb (Supplementary Table S2). In contexts where short read sequencing fails to identify causal variants, a clear vision of the structural variation backbone provided by long read sequencing constitutes a substantial improvement for research and clinical applications.

In a complementary manner, ONT and PacBio yielded by far the best results in haplotype phasing. Phasing has been rarely exploited in past genome profiling studies, because haplotypes remain difficult to phase with short reads without parent-child-trio study design. The advantage of long reads in phasing brings new insights for complex identification of causal variants, involving compound heterozygosity and/or haplotype scenarios responsible for pathologies.

Regarding the optimal depth of coverage for genome profiling, we observed that for all sequencing technologies, lower than the standard 30× mean depth of coverage allows satisfactory analysis of SNVs down to 12×, while indel and SV profiling require at least 20×. In contrast, phasing requires full 30× coverage for the best performance. We observed that DeepVariant mostly performs better than Clair3 in small variant calling, while Sniffles2 shows slight improvement over pbsv in SV calling, regardless of the depth of coverage.

To our knowledge, this study provides the first independent evaluation of ICLR technology. ICLR showed excellent performance in SNV profiling, the best performance in indel profiling, and only an average performance in SV profiling as well as in phasing. We identified several drawbacks of this technology. Firstly, ICLR requires high raw sequence depth of over 200×, which represents high sequencing costs, above the costs of PacBio and ONT, and this will likely prevent wide implementation of ICLR in spite of its performance. Secondly, ICLR requires a bioinformatics treatment in DRAGEN Cloud, with several limitations: it is inflexible in terms of parameter customization; every analysis and reanalysis is payable; and lastly, the algorithm is not available in local instances of DRAGEN, possibly raising issues for certain data-sharing policies.

Sequencing technologies are advancing at an unprecedented pace, and this study benchmarks the current state-of-the-art. Yet, even as we evaluate today’s leading platforms, certain innovations are already emerging that are not yet commercially available. For example, PacBio now provides a new SPRQ Nx chemistry in beta testing (not available during the experimental part of this study), which should further improve the throughput and cost-per-genome without any impact on data quality. Illumina recently released a new Constellation chemistry for beta testing, which might replace ICLR as their reconstructed long read approach; the performance and pricing of the Constellation chemistry are currently unknown. Likewise, the Axelios instrument and associated SBX chemistry from Roche [79], while not commercially available at the time of writing, may require further evaluation in the future.

From a methodological perspective and regarding further implications, long read sequencing is progressively transforming the current gold standard of whole genome sequencing. Related guidelines for bioinformatics analysis are likewise evolving. Long reads are essential for the assembly of more complete reference genomes, with new insights into genomic research provided by the Telomere-to-Telomere consortium [80]. Additionally, recent development of population-specific reference genomes with long reads empowers pangenome analysis, providing better accuracy for genome profiling from both short and long reads using graph genome approaches [13].

This work illustrates that long read technologies provide better mapping, thus enabling haplotype resolution in complex regions, including repeated sequences in various scenarios (pseudogenes, paralogs, transposable elements, etc.). An example of a complex region mapped with short and long read technologies is shown in Supplementary Fig. S6. On this basis, a browsing of the ClinVar database commonly used for targeted regions or gene panel analysis shows that approximately 120k known variants are located in challenging genes referenced in the CMRG set. These variants include more than 9k short indels and 600 insertions/deletions above 50 bp long. Regarding SVs, the ongoing effort to reference SVs with related clinical phenotypes (dbVar ID: nstd102) has so far identified about 84k relevant regions associated to a similar number of known SVs, which would benefit from long read information. While software options for exploiting short read information in challenging genes exist [81–83], long read approaches remain the most efficient solution for *de novo* events characterization, notably for SVs. This point can be illustrated by the rapid development of telomere and centromere analysis toward clinical research application [84, 85].

Taken together, these observations emphasize the pivotal role of long reads in advancing clinical research, leveraging the inclusion of genomic regions previously understudied or disregarded for their complexity. The integration of long reads will provide a more complete understanding of the genome, paving the way for improved diagnosis and therapeutic insights.

## 5 Conclusions

Sequencing technologies are advancing at an unprecedented pace. Using the latest guidelines for bioinformatics processing and benchmarking, we provide a comprehensive evaluation of cutting-edge sequencing technologies. With insight into the pros and cons of different technologies, this study aims to help the decision-making in future research projects, technological platforms development, and clinical applications.

All technologies performed well in SNV calling. These results alleviate existing concerns about missing clinically or biologically significant variants when relying solely on long read technologies. Indel calling was accurate with all technologies except ONT, which lags behind other technologies in spite of the latest improvements in ONT chemistry. In SV calling, both PacBio and ONT long reads showed a clear advantage over short read technologies. Looking specifically at SVs in complex genomic regions (CMRG), ONT achieved better accuracy than PacBio at lower depth of coverage. Additionally, long read technologies achieved a substantially better haplotype phasing compared to short reads.

In conclusion, while short reads remain a valuable option where the cost and throughput are the most critical, long read approaches appear to provide the most complete view of genome variation. Consequently, long read technologies are starting to be integrated into clinical applications and will continue to provide important contributions to genomics research in the coming years.

## Declarations

### Ethics approval and consent to participate

The HG002 sample, part of the Genome in a Bottle (GIAB) consortium, is derived from a publicly available reference genome (NIST Reference Material 8392), intended solely for benchmarking and research purposes.

### Consent for publication

Not applicable

### Availability of data and materials

All raw sequencing data generated in this study at 30× depth of coverage have been submitted to the European Nucleotide Archive (EMBL-EBI) under accession number PRJEB87691 (https://www.ncbi.nlm.nih.gov/bioproject/PRJEB87691). All variant calling files for all depths of coverage generated in this study have been submitted to Zenodo: DOI 15212389 (https://zenodo.org/records/15212389?preview=1&token=eyJhbGciOiJIUzUxMiJ9.eyJpZCI6IjI5MDAxNTZlLTFhMzgtNDk2Yi05MmZkLWY2ZTU1MjU4ZWE2NCIsImRhdGEiOnt9LCJyYW5kb20iOiI2ZDc2YjM0M2M5ODllZTUxYmQyNTFkMTRmNDFiYmYzZSJ9.HCUhlZD9eYxly6ZC3uPIZQ92En8Vmvc88G9F8X1hODNbBLxRqAI5nEHbRnt5W01tMogKyjSEkEmlOJXekK8LcQ).

### Authors’ contributions

VM, ZG, AL, and AB conceived the study. MD, CF, CD, and ZG performed the experiments. AL and AB performed data analyses and generated figures, with input from FS, DD, SM, and VM. VM, ZG, and JFD supervised the project. DBD, FR, and RO supervised data production. RO, JFD, and VT secured funding for the project. VM, ZG, AL, AB wrote the manuscript with input from FS and DD. All authors reviewed the manuscript.

## Acknowledgements

We would like to express our gratitude to all members of the CNRGH who contributed to this study, particularly from the following labs: LB, L2PGH, LD, LIMS, and LBI.

## Supplementary Data

### 1 Bioinformatic workflow

#### 1.1 Mapping

**Supplementary Template 1 —** Command lines for the alignment process.

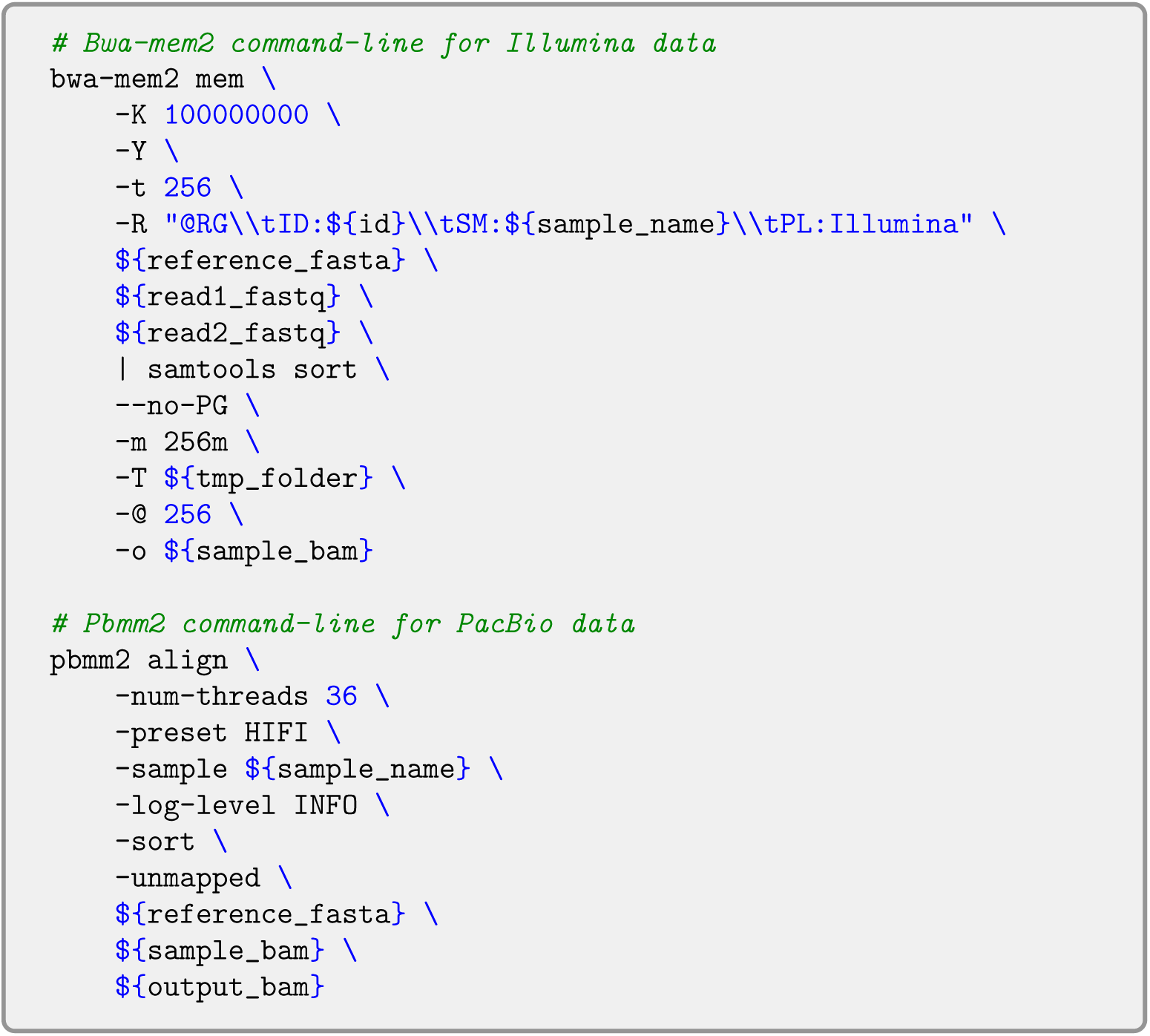

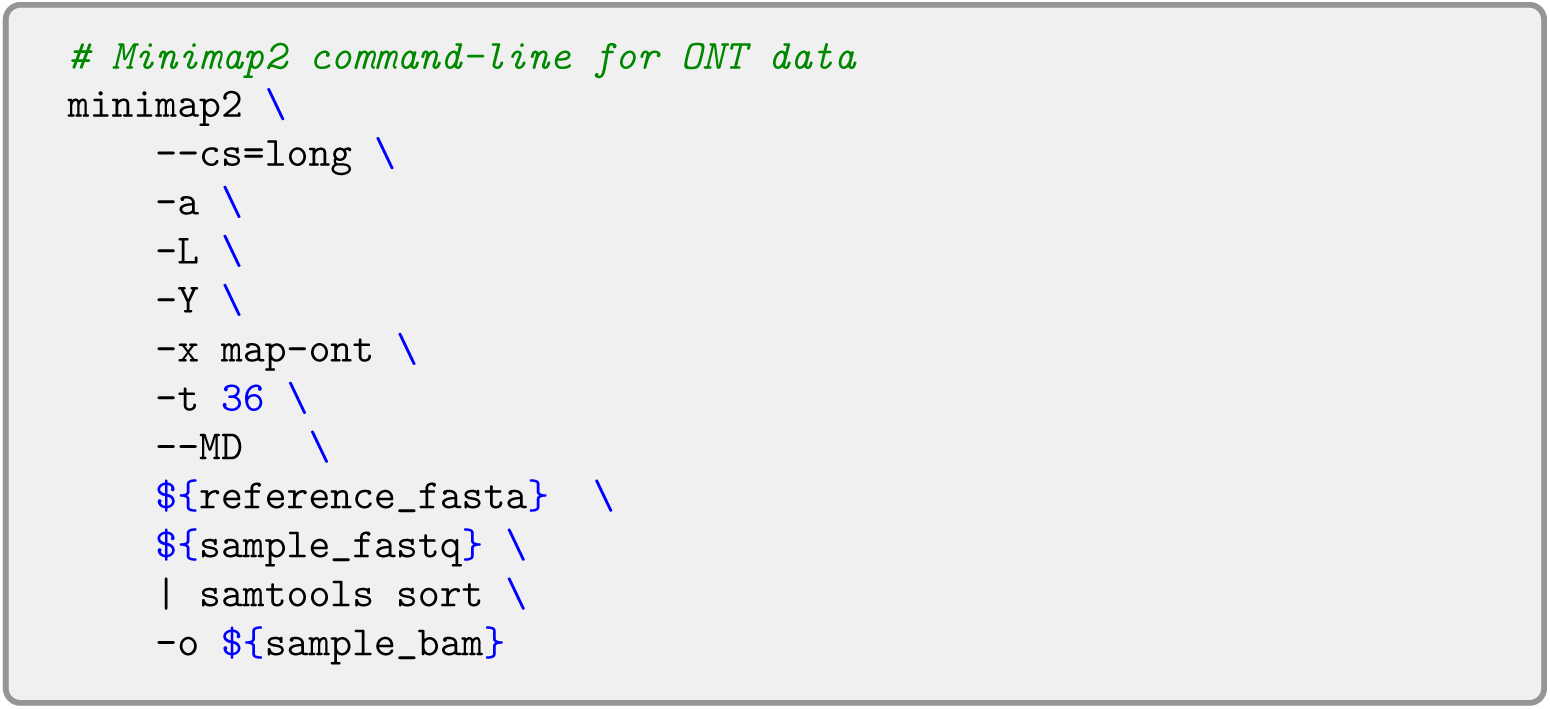

#### 1.2 Downsampling

**Supplementary Template 2 —** Command lines for the downsampling step.

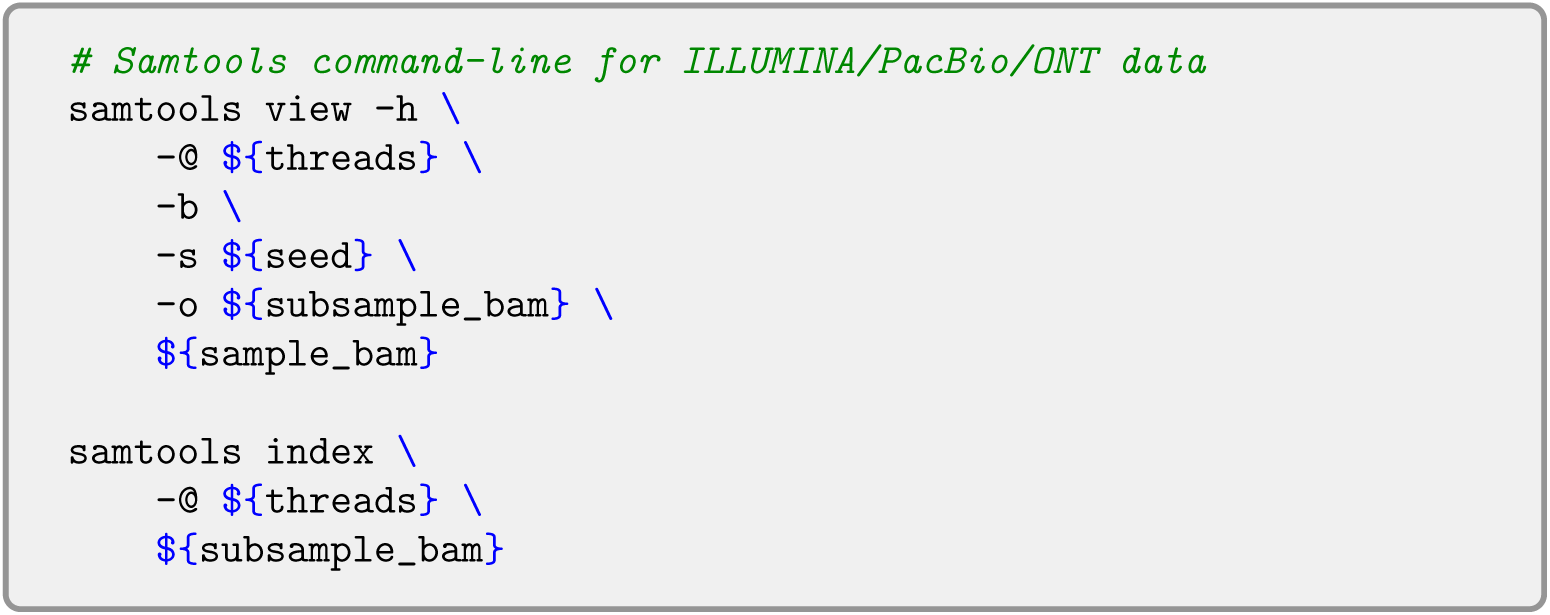

#### 1.3 Structural variation calling

**Supplementary Template 3 —** Command lines for structural variant calling.

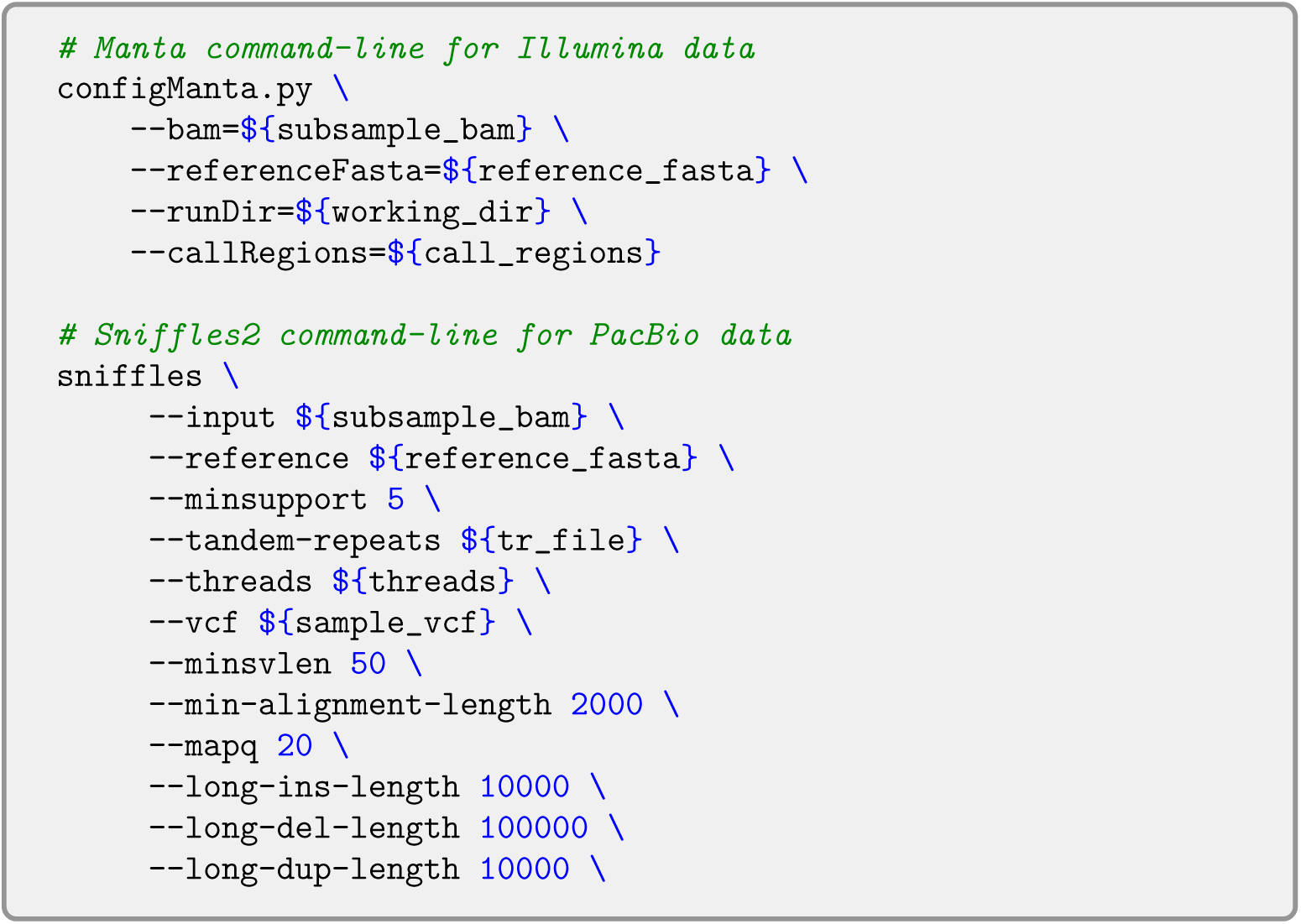

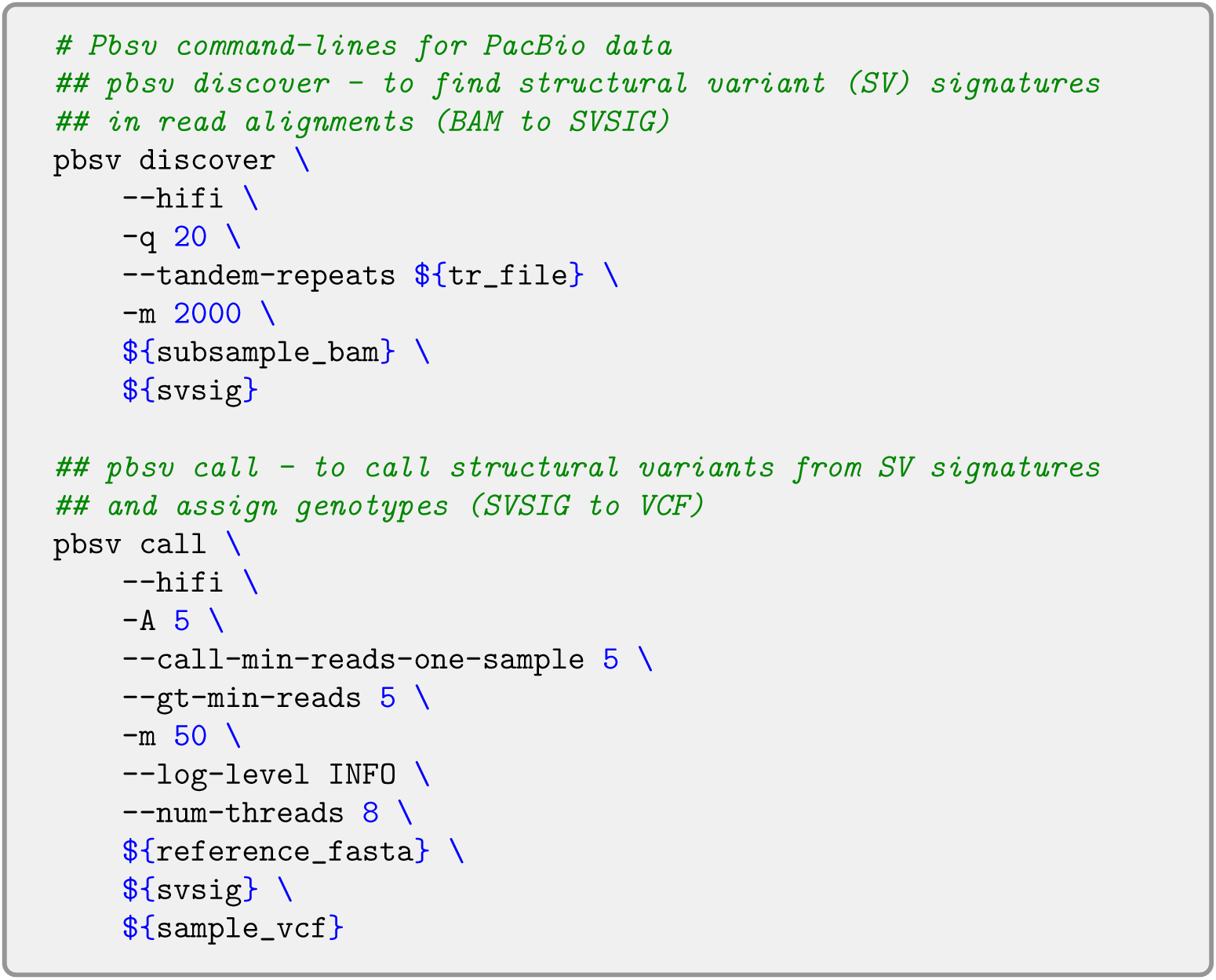

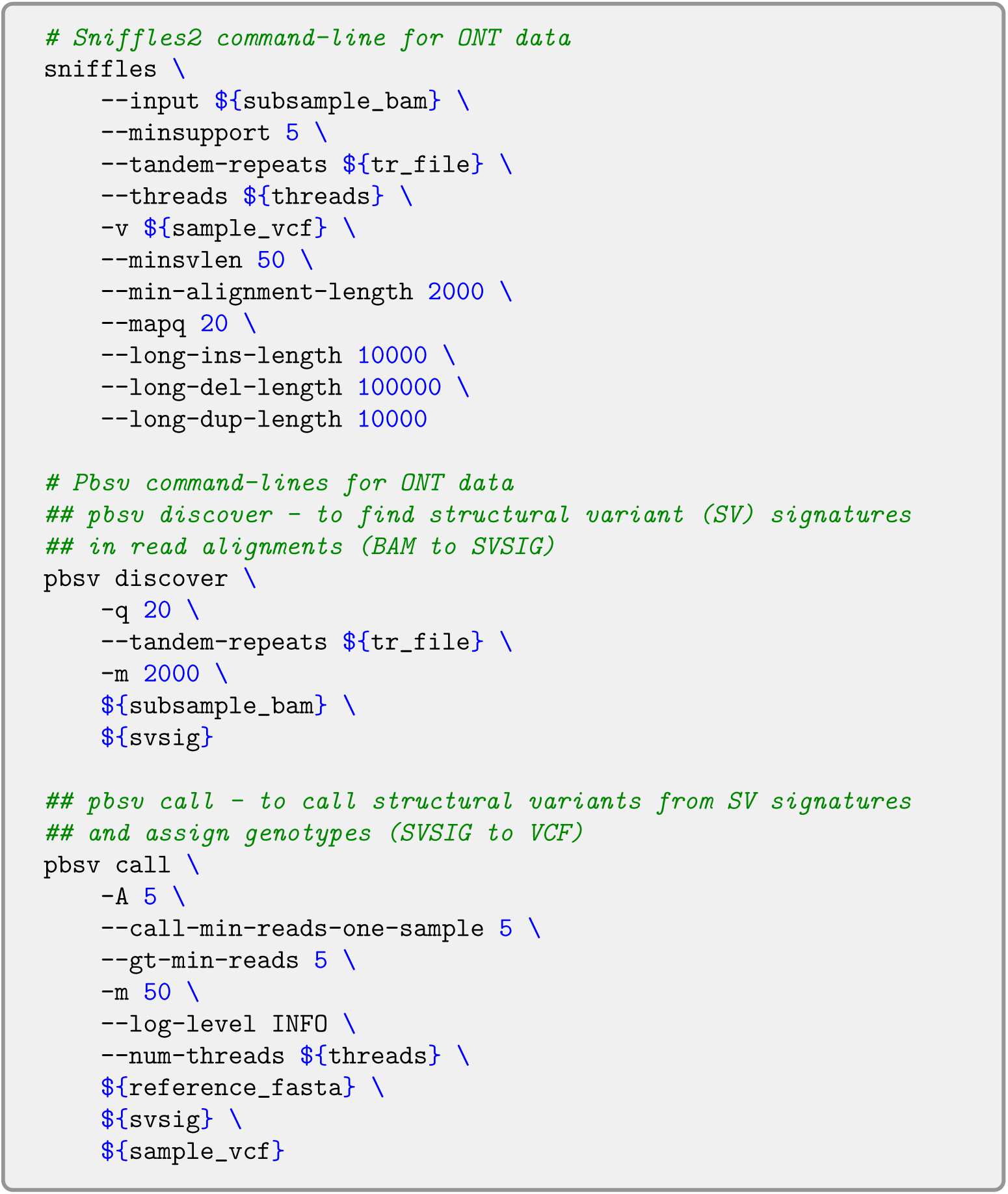

#### 1.4 Small variants calling

**Supplementary Template 4 —** Command lines for small variant calling.

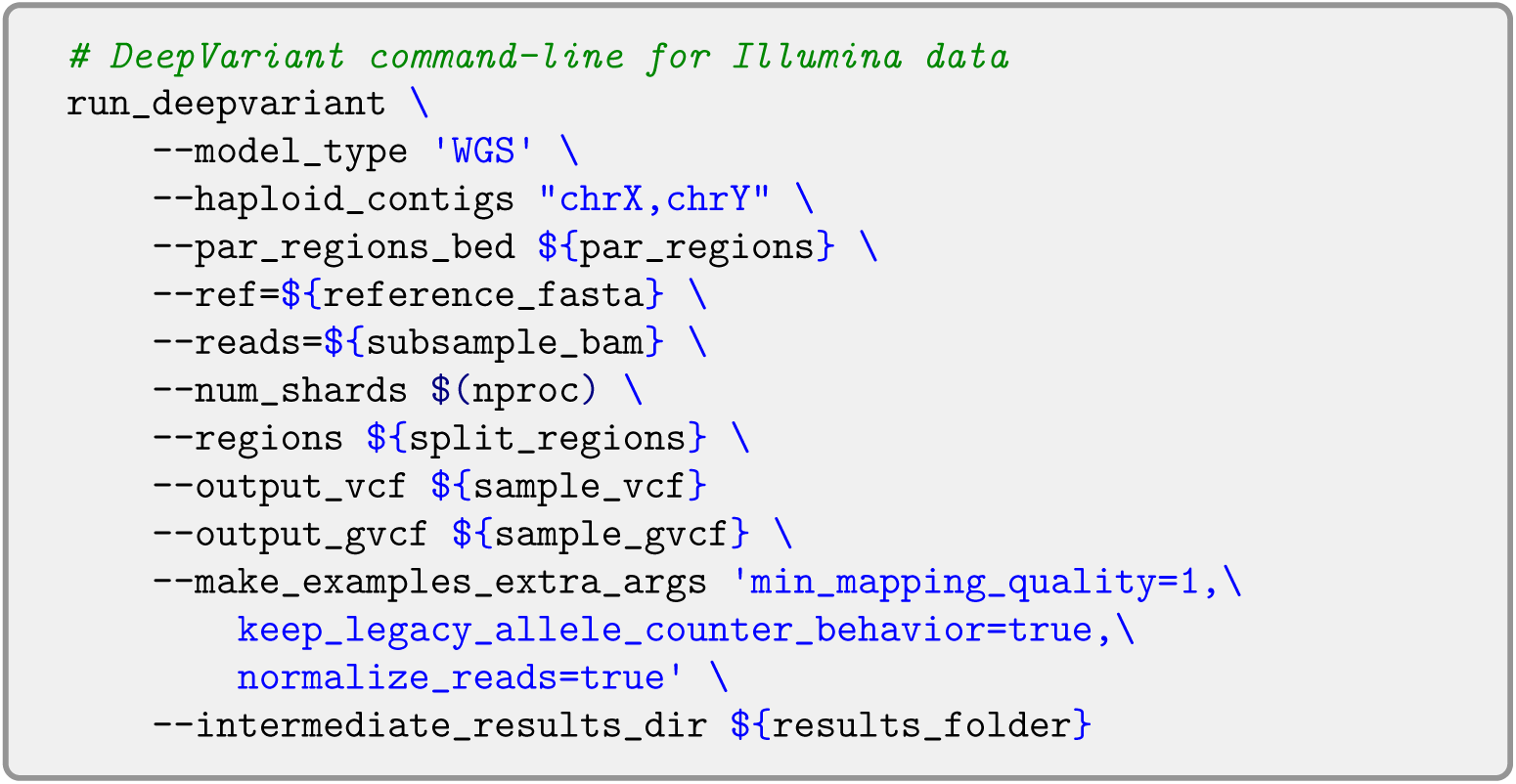

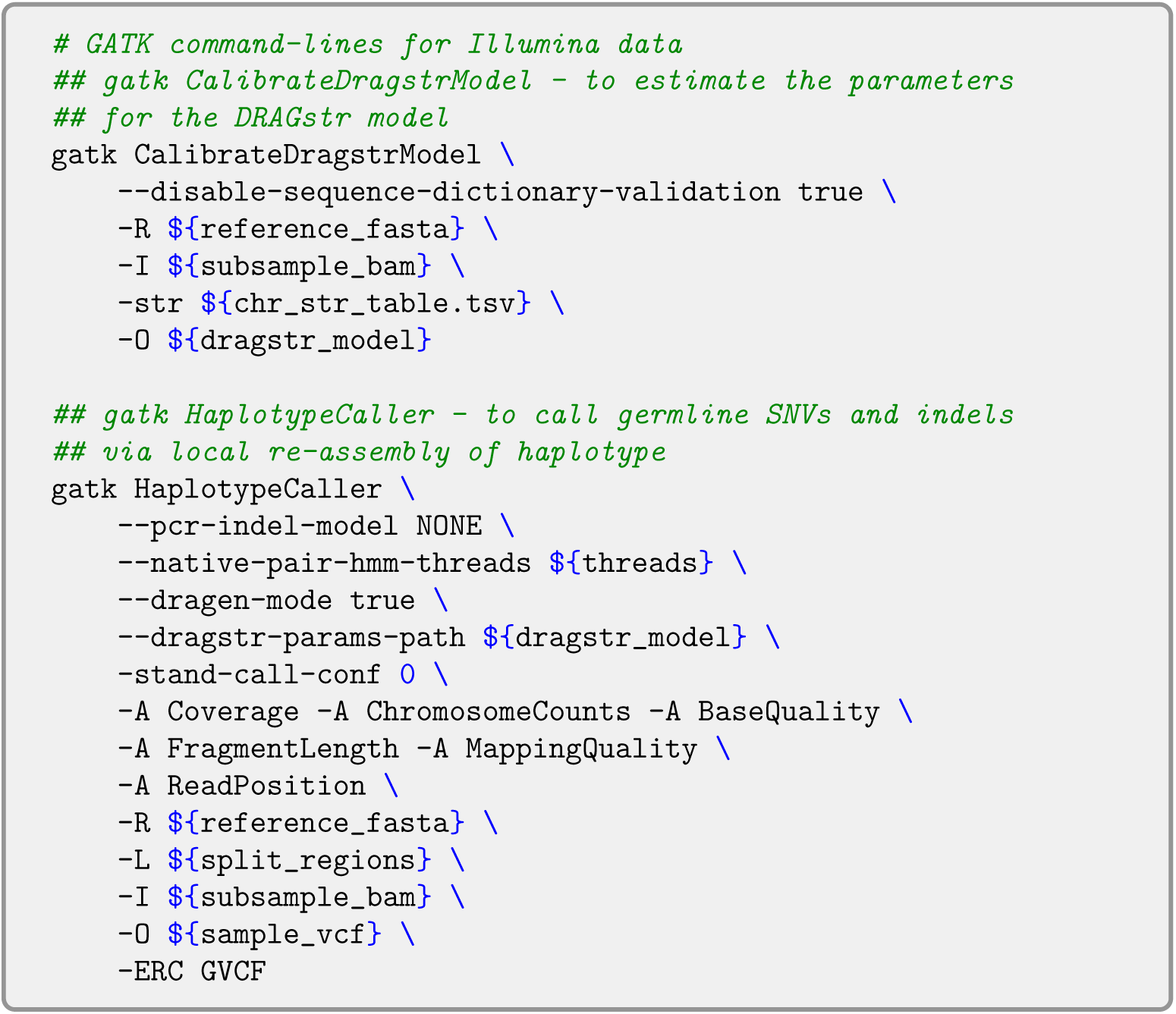

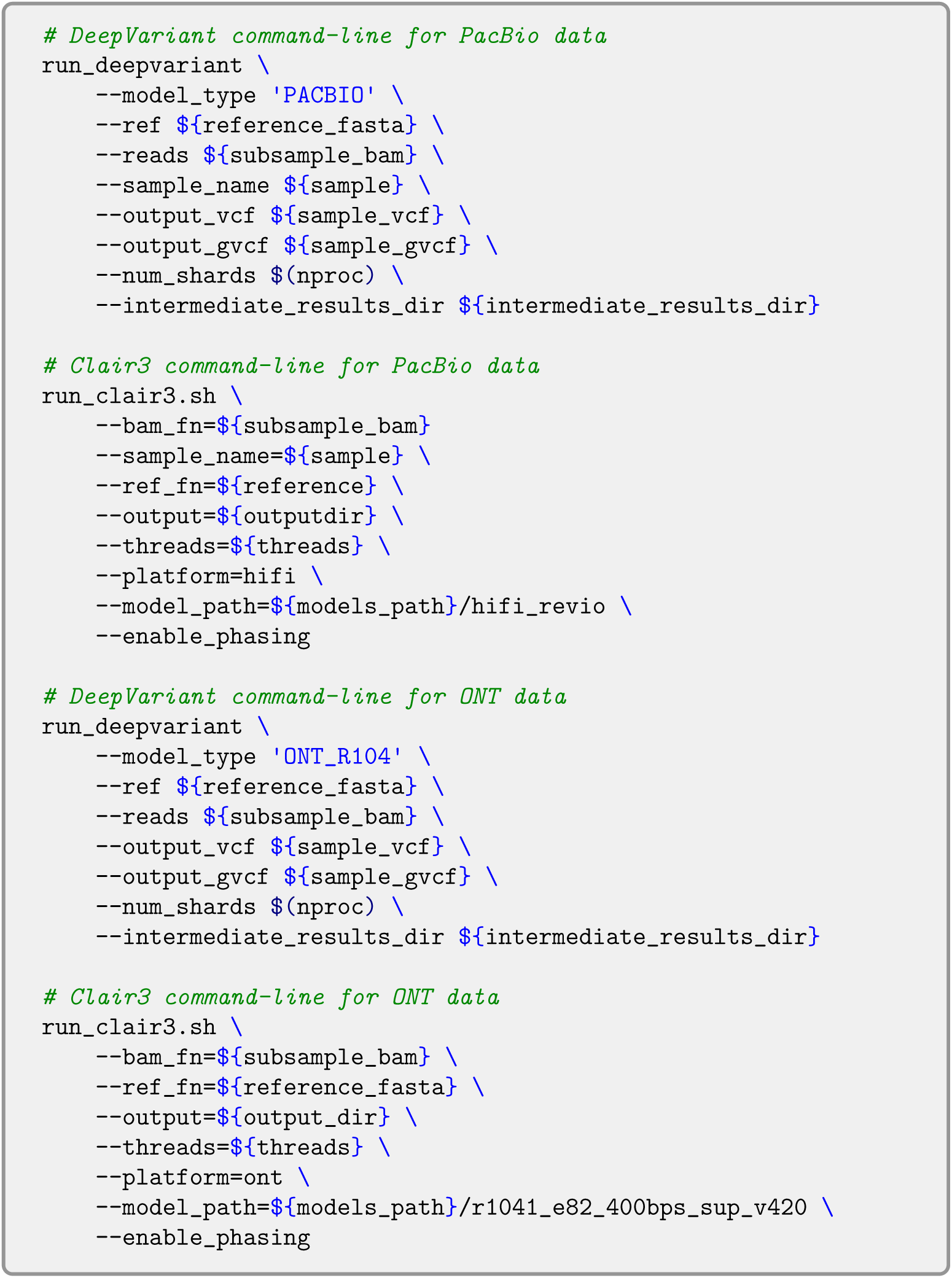

#### 1.5 Phasing

**Supplementary Template 5 —** Command lines for the haplotype phasing.

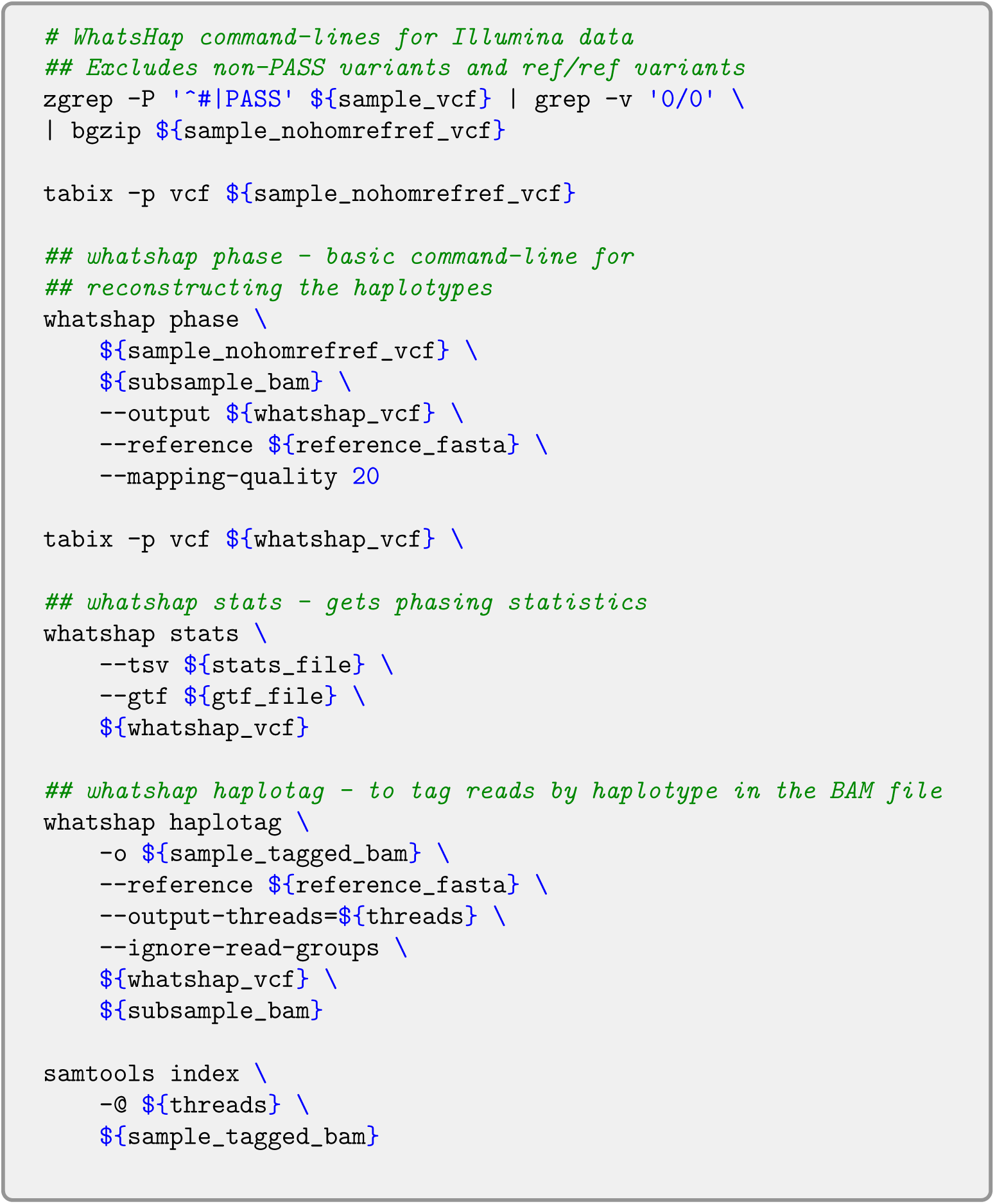

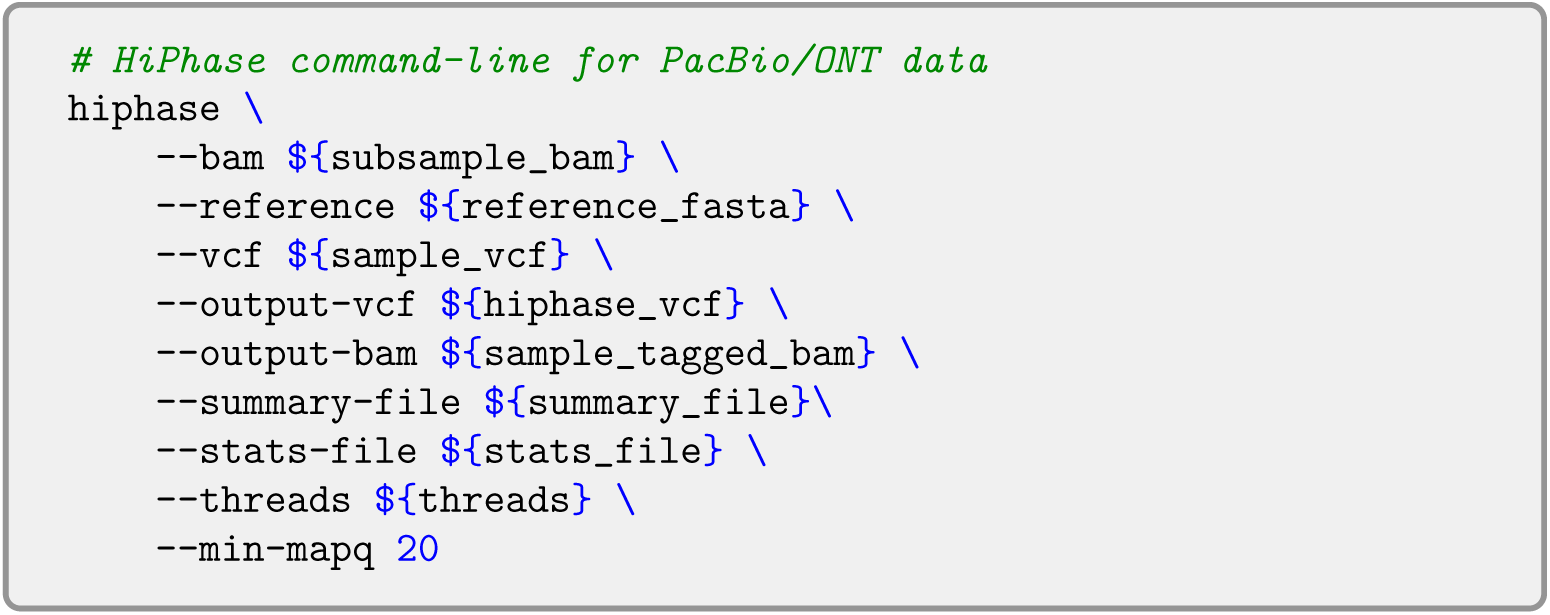

#### 1.6 Assessment of structural variant calling

**Supplementary Template 6 —** Command line for the assessment of structural variant calling with Truvari bench.

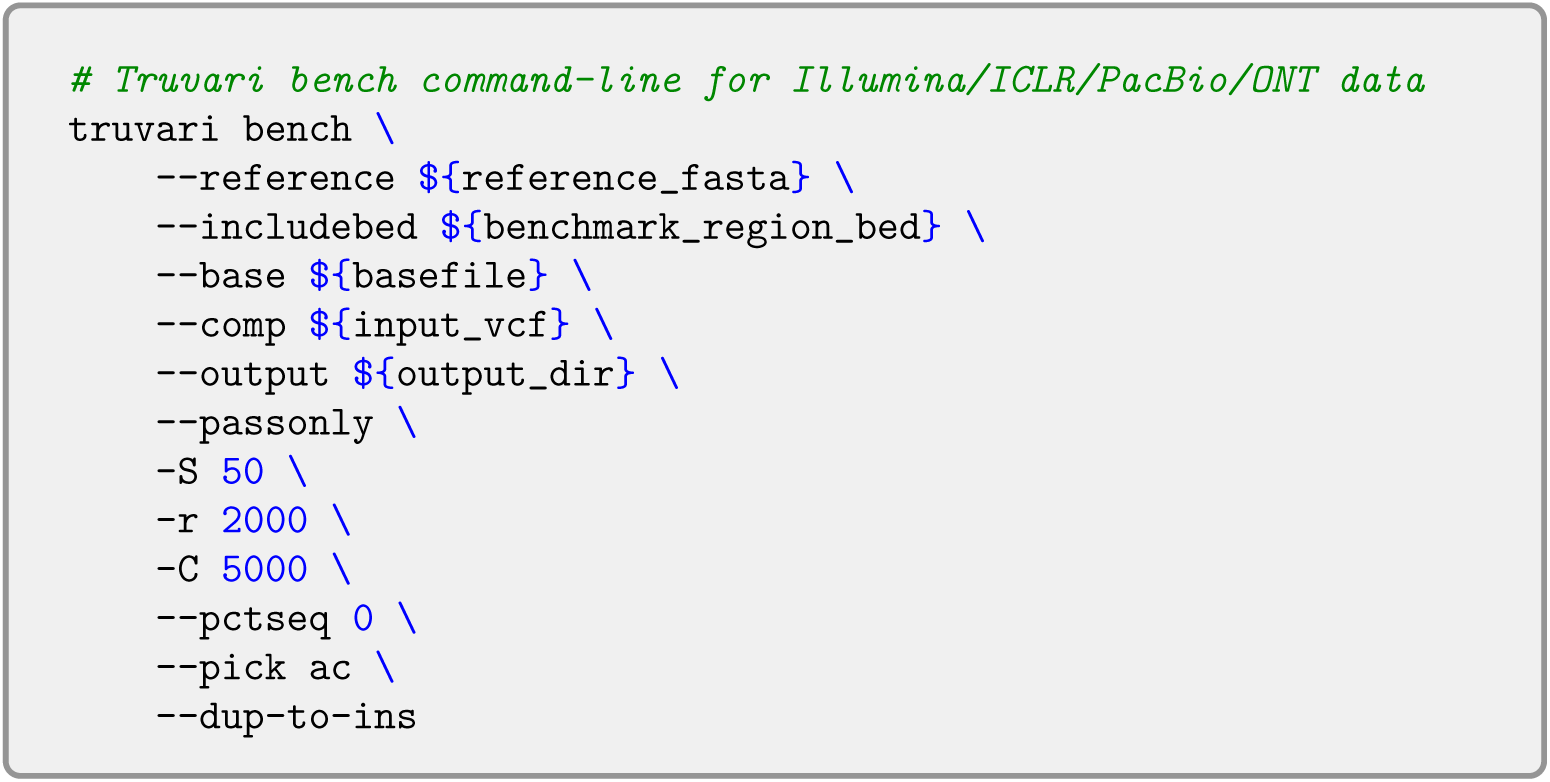

**Supplementary Template 7 —** Command lines for the assessment of structural variant calling with Truvari refine and ga4gh.

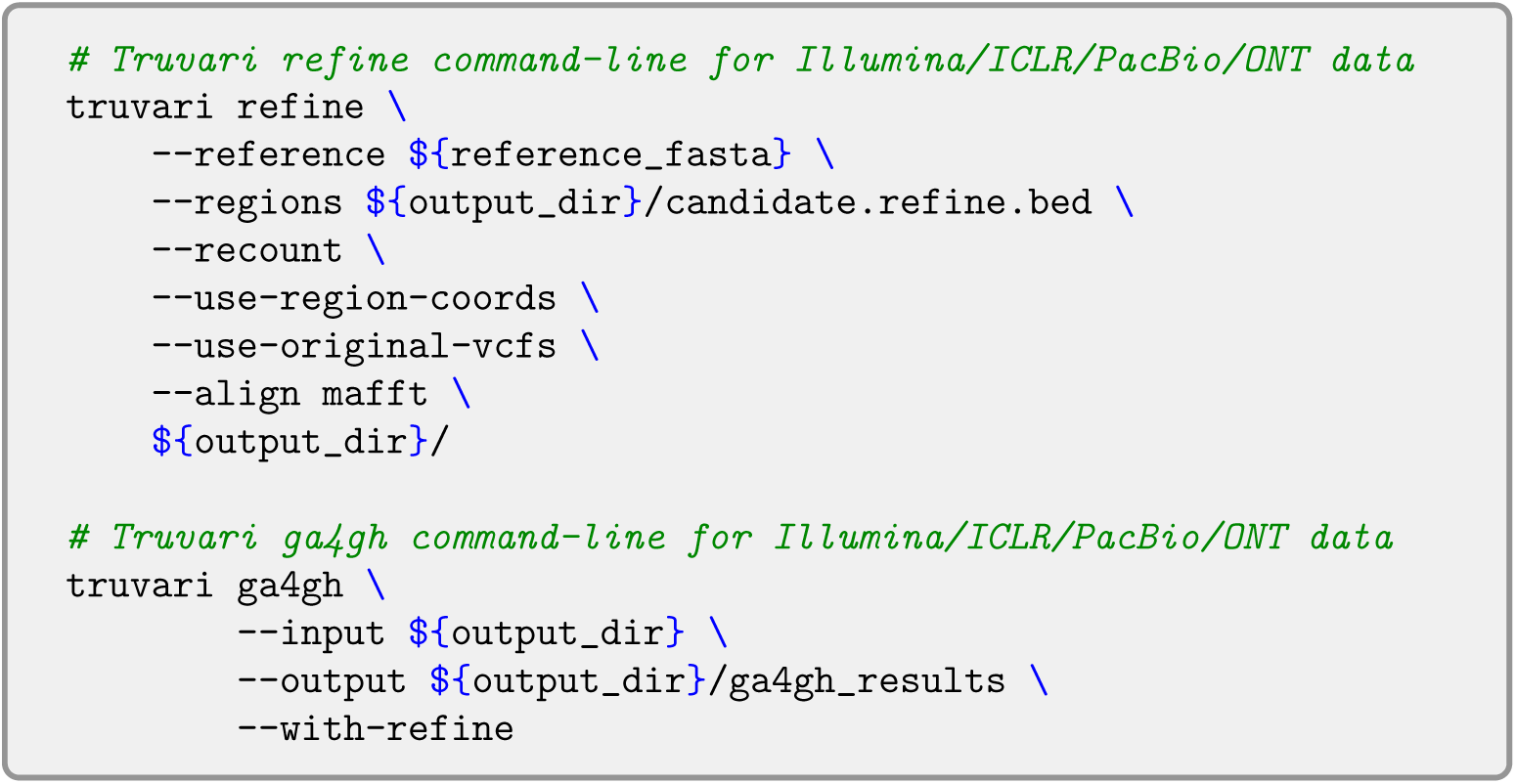

#### 1.7 Assessment of small variant calling

**Supplementary Template 8 —** Command lines for the assessment of small variant calling.

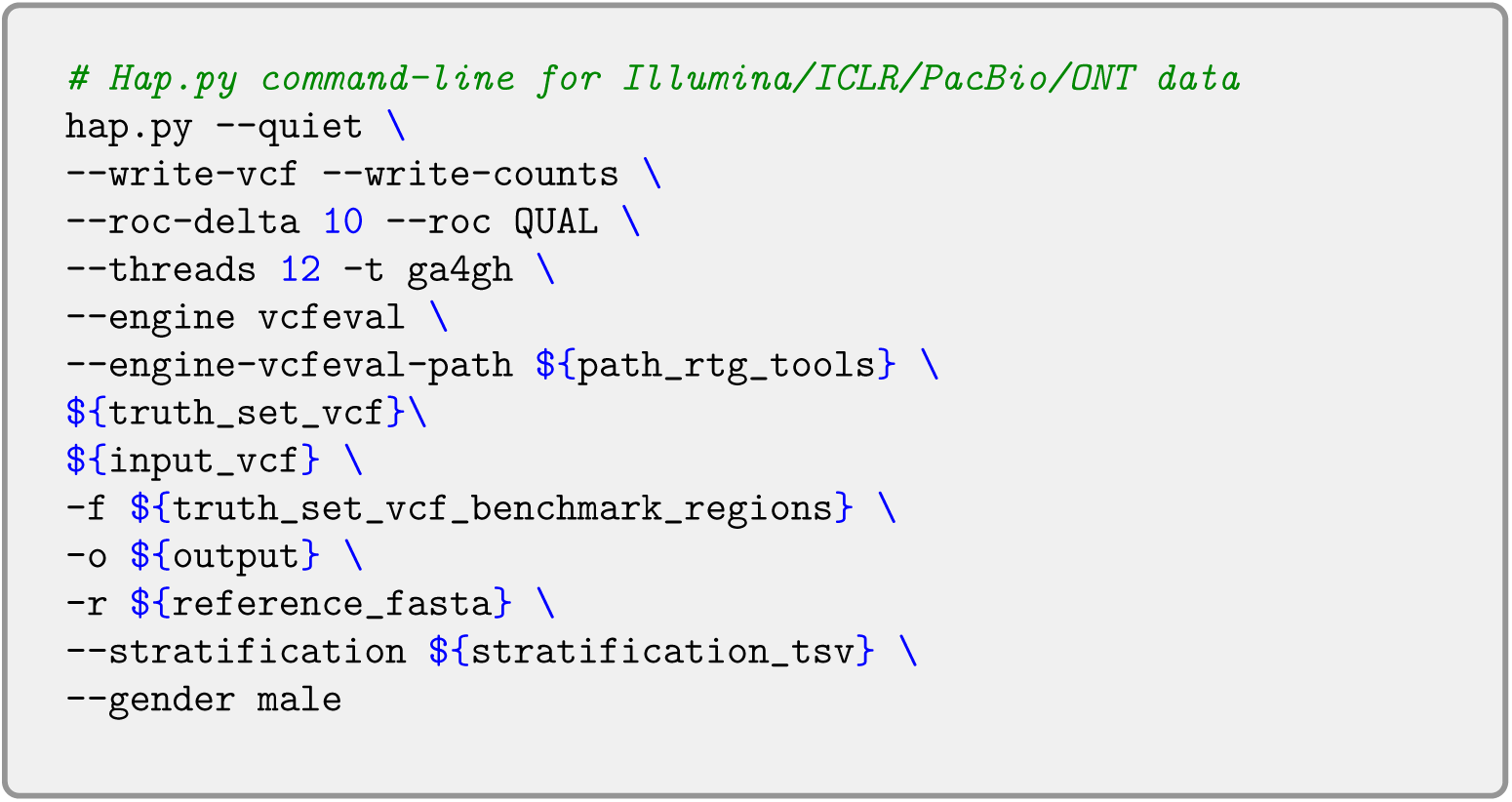

#### 1.8 Phasing assessment

**Supplementary Template 9 —** Command lines for the assessment of phasing.

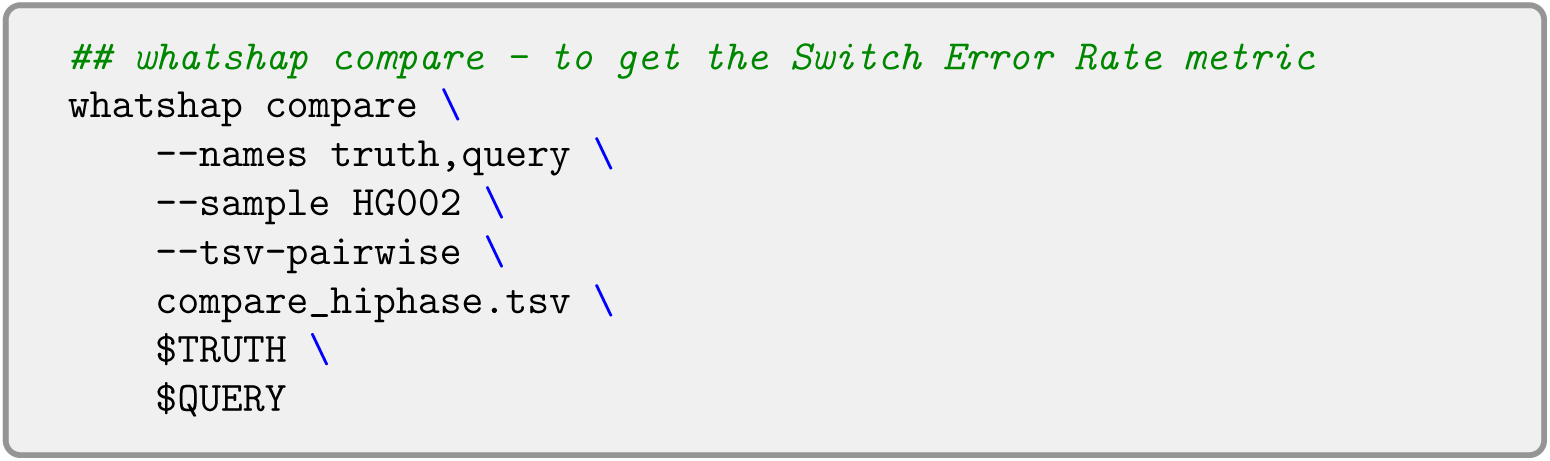

### 2 Truth sets

The Genome in a Bottle Telomere-to-Telomere (GIAB T2T) [66], Challenging Medically Relevant Genes (CMRG) [68], and The Human Genome Structural Variation Consortium (HGSVC) [70] truth sets are high-confidence benchmark datasets designed to improve structural and small variant detection in complex genomic regions.

GIAB T2T-HG002-Q100v1.1 is a telomere-to-telomere (T2T) reference dataset for HG002 obtained by a diploid assembly aligned to GRCh38, offering a nearly complete, gapless human genome to enhance benchmarking accuracy. CMRG truth set targets clinically significant genes in difficult genomic regions, providing a high-confidence benchmark for medical genetics and precision medicine. HGSVC Freeze 4 focuses on diverse human genomes, incorporating phased structural variants from long read sequencing to capture population-level variation. Together, these datasets support the evaluation of sequencing technologies and variant calling tools, ensuring more accurate genomic analyses for both research and clinical applications.

**Supplementary Table S1.** The total number of structural variants (SVs) and small variants in the truth sets used in this study: Genome in a Bottle Telomere-to-Telomere (GIAB T2T), Challenging Medically Relevant Genes (CMRG), and The Human Genome Structural Variation Consortium (HGSVC).

|  | SV |  | Small Variants |  |
| --- | --- | --- | --- | --- |
| | INS ($\geq 50$bp) | DEL ($\geq 50$bp) | SNV | Indels |
| GIAB T2T | 21 133 | 13 118 | 3 461 568 | 586 728 |
| CMRG | 125 | 95 | 21 510 | 6 427 |
| HGSVC | 14 012 | 9 038 | NA | NA |

**Supplementary Fig. S1.**
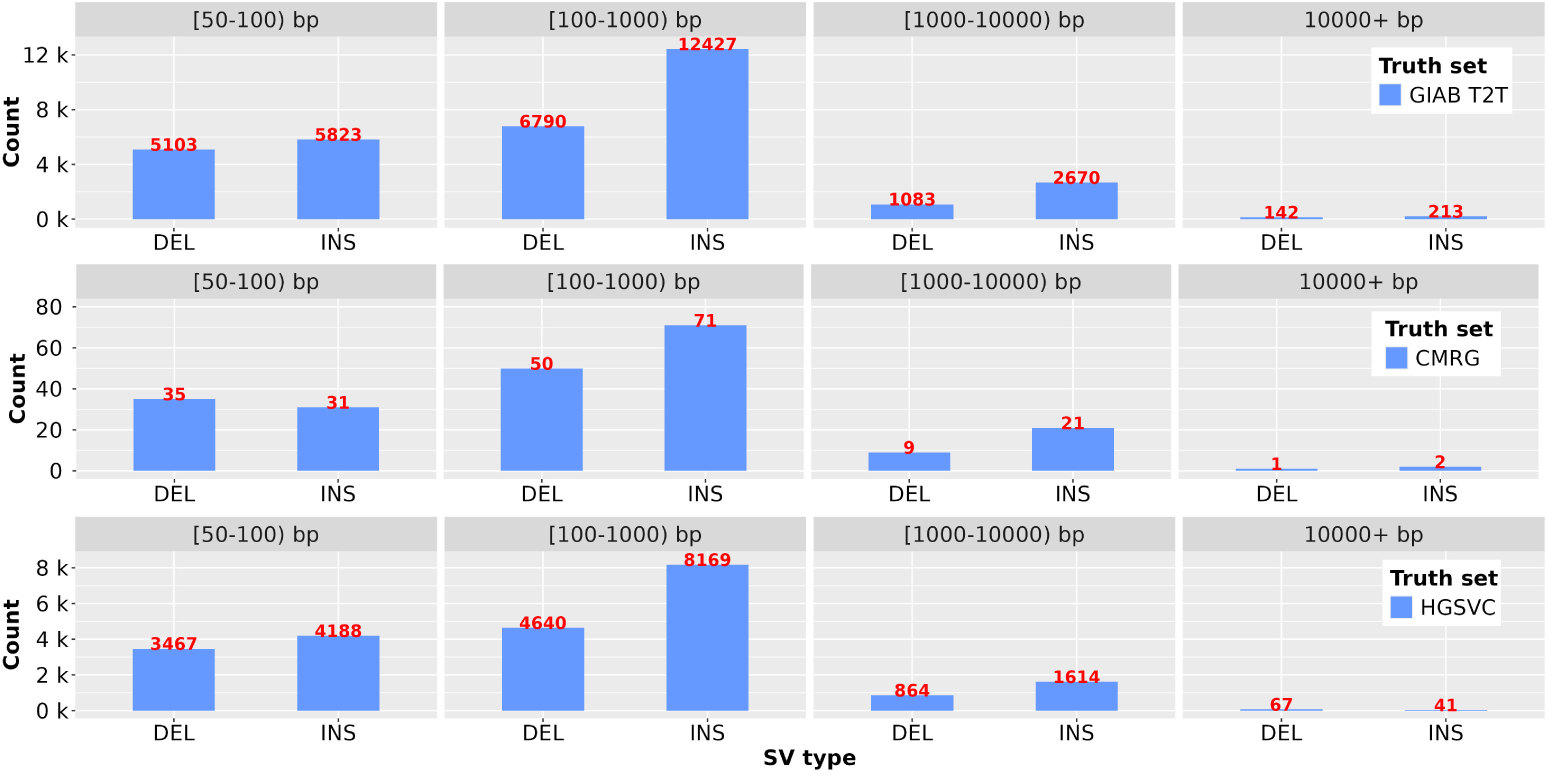
The number of structural variants (SVs) in the truth sets used in this study: Genome in a Bottle Telomere-to-Telomere (GIAB T2T), Challenging Medically Relevant Genes (CMRG), and The Human Genome Structural Variation Consortium (HGSVC). The SVs are categorized into deletions (DEL) and insertions (INS) divided by size into four classes : [50-100) bp, [100-1000) bp, [1000-10000) bp, and 10000+ bp.

### 3 Evaluation metrics

The performance of variant detection was assessed using three key metrics: Precision, Recall, and F1 score.

#### Precision

Precision is defined as the proportion of true positive predictions among all predicted positives:

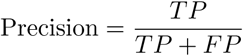

where *TP* is the number of true positives and *FP* is the number of false positives.

#### Recall

Recall (also known as Sensitivity) is the proportion of true positive predictions among all actual positives:

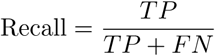

where *TP* is the number of true positives and *FN* is the number of false negatives.

#### F1 score

The F1 score is the harmonic mean of Precision and Recall, providing a balanced measure of both:

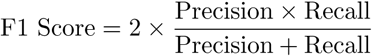

The F1 score ranges from 0 to 1, with 1 being the best possible performance.

### 4 Structural variant calling

Supplementary Fig. S2 illustrates the quantity of various identified SV types detected at 30× coverage across all sequencing technologies: Illumina short reads (Illumina), Illumina Complete Long Reads (ICLR), Pacific Biosciences (PacBio), Oxford Nanopore Technologies (ONT). The SV caller Sniffles2 was assessed here for PacBio data.

**Supplementary Fig. S2.**
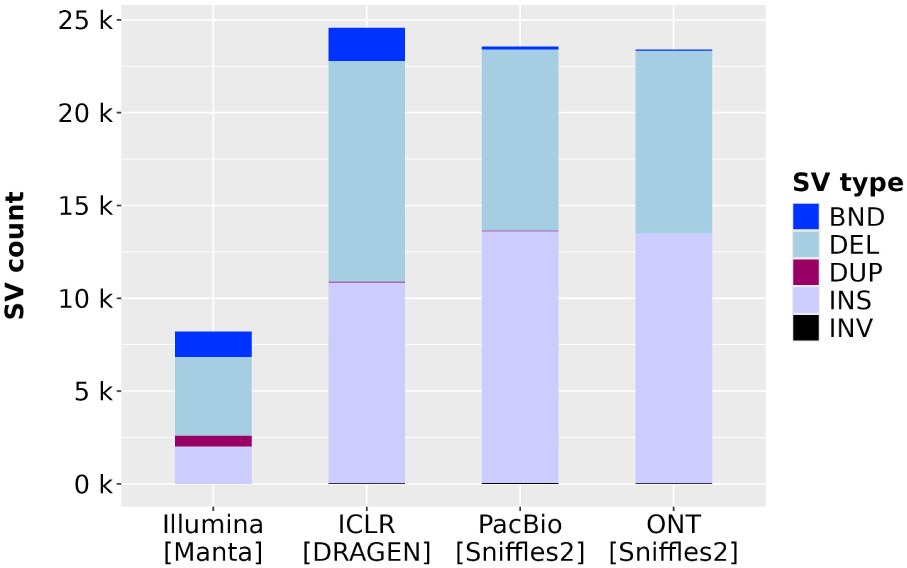
The count of different structural variant (SV) types detected at 30× across all sequencing technologies. BND - breakend events, DEL - deletions, DUP - duplications, INS - insertions, INV - inversions.

**Supplementary Fig. S3.**
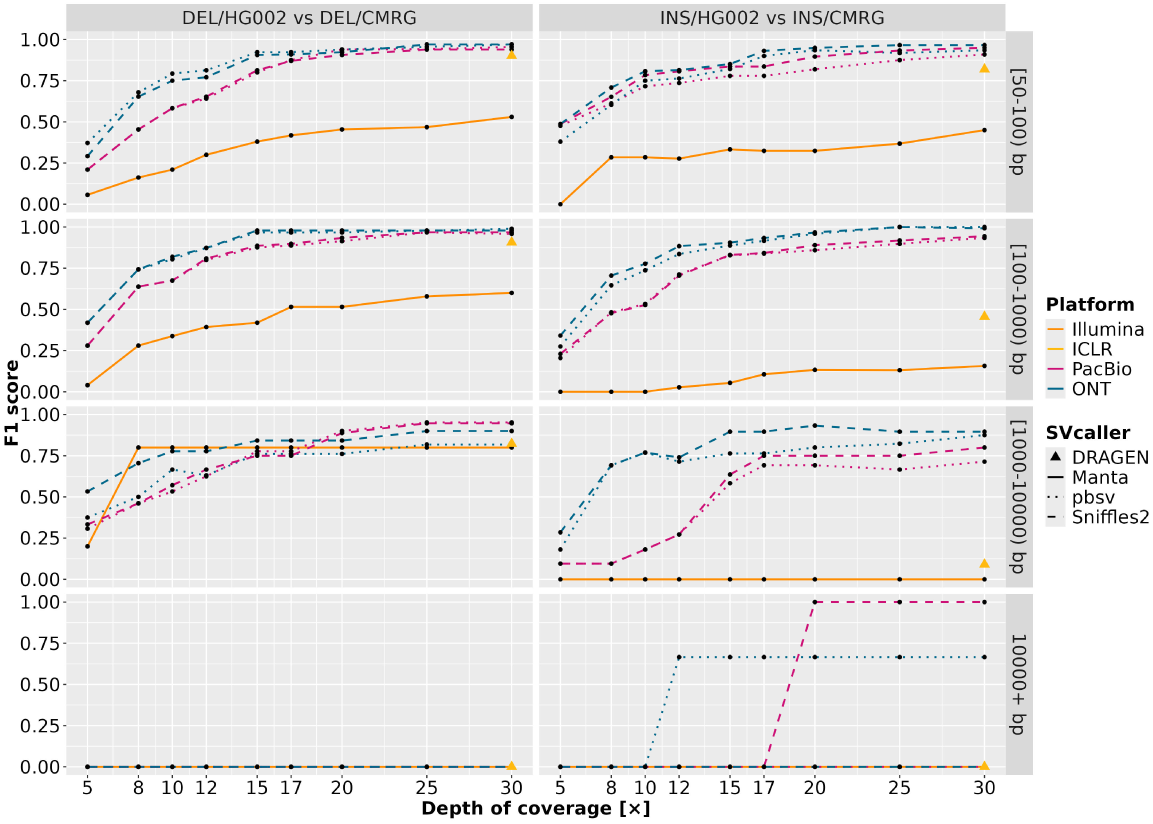
The decline of F1 score with decreasing mean depth of coverage for deletions (DEL) and insertions (INS) for different size classes of structural variants, compared to the Challenging Medically Relevant Genes (CMRG) truth set.

**Supplementary Fig. S4.**
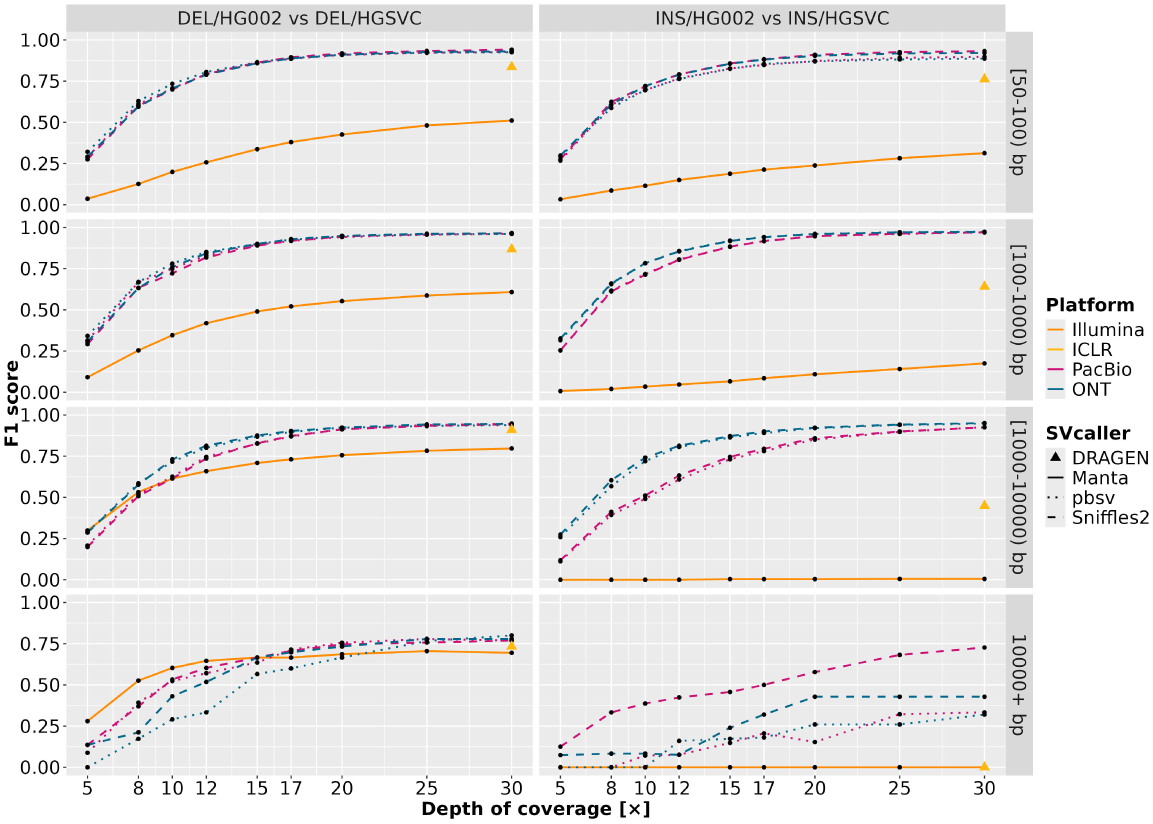
The decline of F1 score with decreasing mean depth of coverage for deletions (DEL) and insertions (INS) for different size classes of structural variants, compared to The Human Genome Structural Variation Consortium (HGSVC) truth set.

**Supplementary Table S2.** The number of SVs and related cumulative size within the comparison of Illumina short reads against ONT and PacBio. After comparison against the GIAB T2T high-confidence truth set, the Illumina true positives were compared to ONT (Sniffles2) and PacBio (pbsv) true positives as reference datasets, respectively, using the “truvari bench” program and parameters described below in Section 1.6. The SV count for each output was computed with the “bcftools stats” program. The cumulative size was determined from available “SVLEN” tags within the ONT and PacBio vcf files. The limited availability of “SVLEN” information provided by Manta and Illumina short reads does not allow the computation of a relevant SVs cumulative size.

| Technologies | Comparison status | SVs count | Cumulative size (bp) |
| --- | --- | --- | --- |
| Illumina vs PacBio | Shared GIAB T2T high-confidence truth set SVs | 4 613 | 2 570 691 |
| Illumina vs PacBio | PacBio-specific GIAB T2T high-confidence truth set SVs | 13 965 | 4 374 544 |
| Illumina vs PacBio | Illumina-specific GIAB T2T high-confidence truth set SVs | 308 | - |
| Illumina vs ONT | Shared GIAB T2T high-confidence truth set SVs | 4 463 | 2 510 097 |
| Illumina vs ONT | ONT-specific GIAB T2T high-confidence truth set SVs | 12 230 | 4 845 839 |
| Illumina vs ONT | Illumina-specific GIAB T2T high-confidence truth set SVs | 458 | - |

### 5 Small variant calling

Supplementary Fig. S5 displays the progression of the F1 score for SNVs and indels compared to the CMRG truth set at different mean depths of coverage among all platforms: Illumina, ICLR, PacBio and ONT.

**Supplementary Fig. S5.**
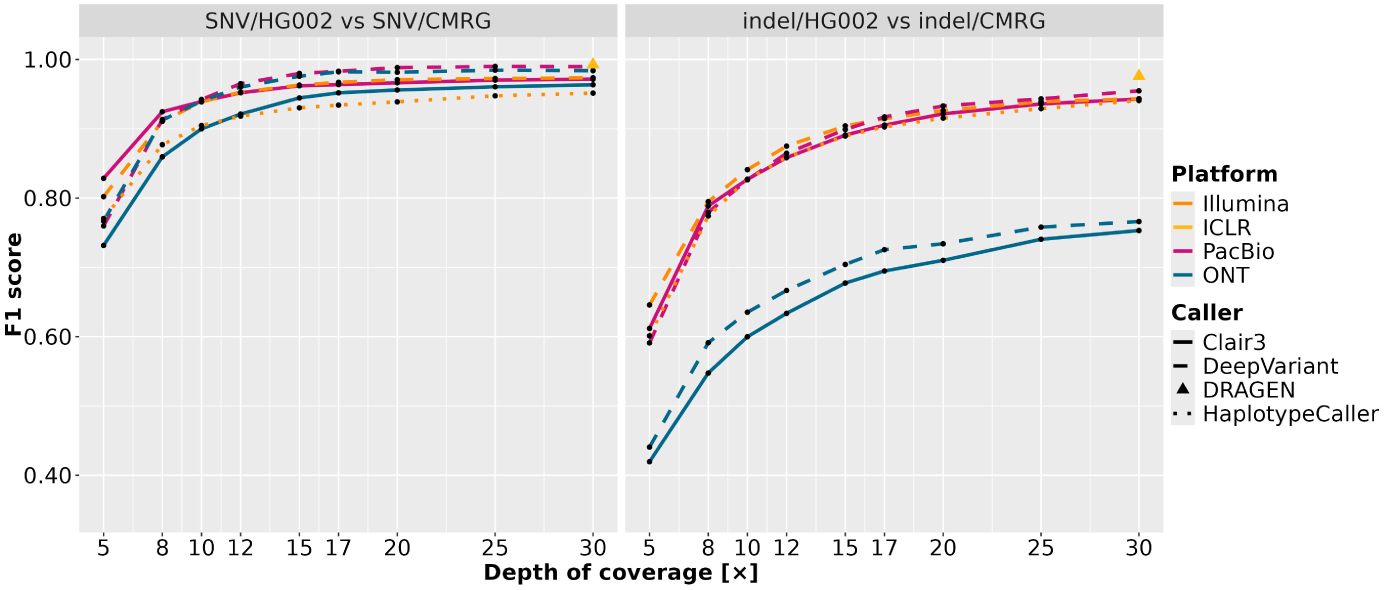
The decline of F1 score with decreasing mean depth of coverage for single nucleotide variants (SNVs) and insertion-deletions*<*50 bp long (indels) for the Challenging Medically Relevant Genes (CMRG) truth set.

### 6 SMN1 gene coverage comparison

Supplementary Fig. S6 shows an IGV [v2.17.1] (Integrative Genomics Viewer [86]) snapshot of the mapped reads organization of the SMN1 genomic region at 30× coverage for Illumina, ICLR, PacBio and ONT (GRCh38).

**Supplementary Fig. S6.**
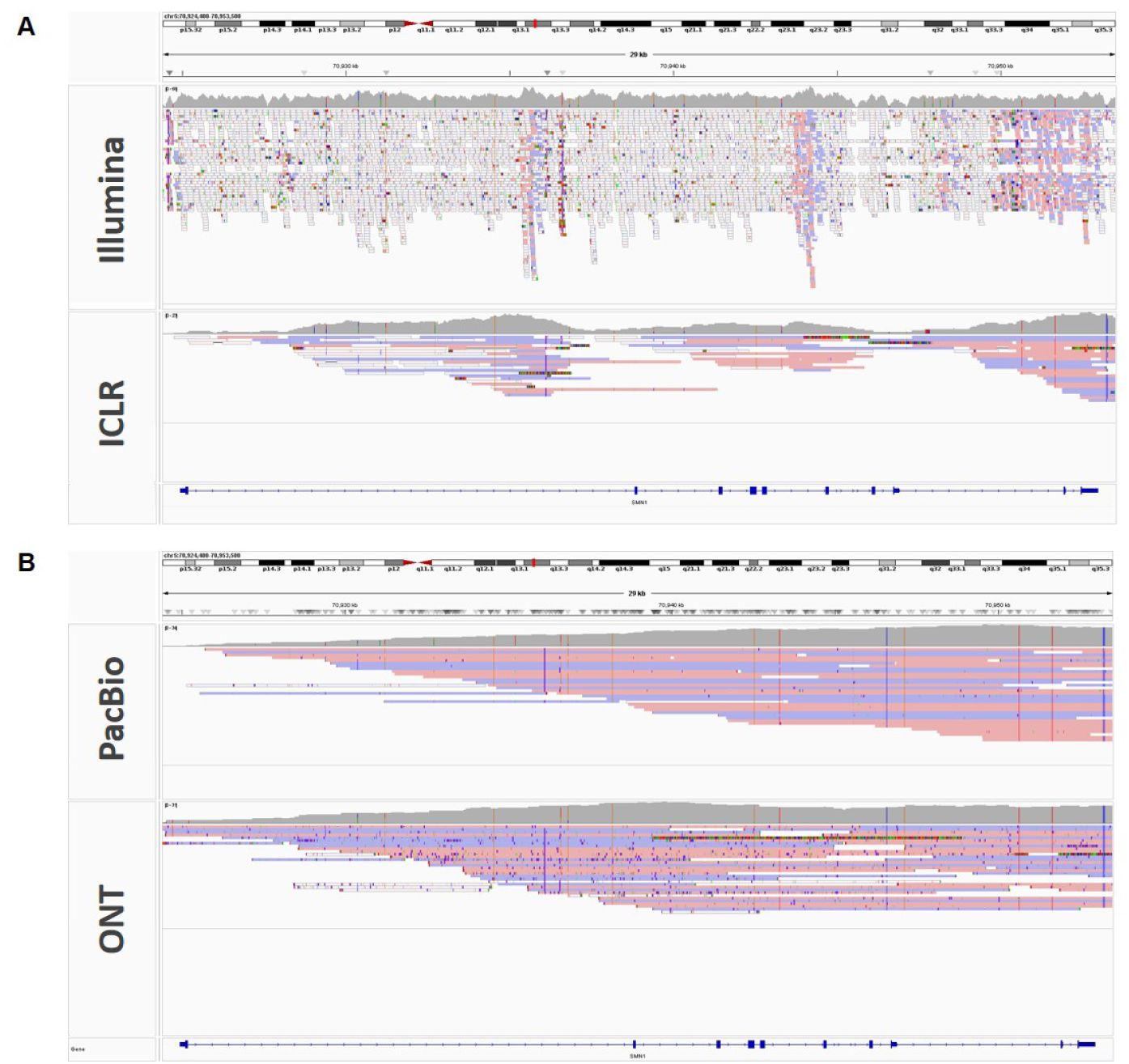
The SMN1 gene coverage organization. **A)** Illumina short reads (Illumina) and Illumina Complete Long Reads (ICLR) **B)** Pacific Biosciences (PacBio) and Oxford Nanopore Technologies (ONT).

